# Phactr4 influences macrophage lamellipodial structure and dynamics through Arp2/3 complex and Ezrin regulation

**DOI:** 10.1101/2025.05.13.653717

**Authors:** Rohini Manickam, Jeremy D. Rotty

## Abstract

Dynamic cycles of actin remodeling drive membrane protrusion and retraction events essential for macrophage function. Phosphoregulation of actin-associated proteins plays a key role, but the factors that determine the spatiotemporal balance between kinases and phosphatases is less well understood in this context. Here, we identify the Protein Phosphatase 1 (PP1)-binding protein Phactr4 as a critical regulator of cytoskeletal remodeling. Phactr4 loss disrupts lamellipodial architecture, which results in uncoordinated migration and disrupted iC3b-mediated phagocytosis. Unstable membrane dynamics underlie the Phactr4 knockdown phenotypes. Phactr4 is recruited to the leading edge via interaction with active Arp2/3 complex, and strongly correlates with membrane retraction. Phactr4 loss leads to ezrin hyperphosphorylation, and membrane protrusion defects in these cells are reversed by ezrin inhibition. Our findings position Phactr4 as a critical PP1-dependent coordinator of cytoskeletal remodeling during macrophage migration and phagocytosis. Recent reports have linked Phactr4 to several human disease states, which may be due to its influence on actin dynamics.

## INTRODUCTION

Dynamic membrane protrusions allow innate immune cells, such as macrophages, to continuously probe the extracellular environment as they perform essential physiological functions like phagocytosis, macropinocytosis and migration [1]. Precise regulation of dynamic actin within phagocytic cups and migratory lamellipodial protrusions is critical for processes like immune surveillance. Cytoskeletal dysfunction impairs pathogen clearance, disrupts inflammatory responses and contributes to immunopathology [2]. Outside of host defense, membrane protrusions respond efficiently to the microenvironment and contribute to tissue surveillance and homeostasis [3]. Despite the importance of the actin cytoskeleton to these processes, its regulation at the molecular level likely remains incompletely understood.

Dynamic cellular protrusions rely on the assembly and disassembly of actin filaments, which is orchestrated by a complex array of actin-associated proteins. The actin-related protein 2/3 (Arp2/3) complex nucleates dense branched actin networks that play a fundamental role in lamellipodial protrusion at the cell’s leading edge [4]. While lamellipodial protrusion is not universally required for all environmental sensing, integrin-dependent motility and phagocytosis require Arp2/3 complex activity to sense and respond to extracellular matrix and complement proteins [5]. Productive membrane extension also requires a balance between actin polymerization and turnover at the leading edge. The ADF/cofilin family of actin severing proteins generates new barbed ends that sustain actin treadmilling and allow for continued protrusion [6]. Significant progress has been made in understanding how cytoskeletal regulators drive membrane protrusions, but how these processes are spatiotemporally coordinated remains an open question.

Phospho-regulation of leading-edge cytoskeletal proteins greatly impacts their activity, and may play a key role in spatiotemporal regulation of lamellipodial protrusion. For example, the activity of Arp2/3 complex, cofilin, coronins, ERM proteins (ezrin, radixin, moesin), among many others, is greatly impacted by specific phosphorylation events. One classic example is Ser3 phosphorylation of cofilin, which lowers its actin severing ability [7]. Hyper-phosphorylation of Ser3 has been linked to defects in lamellipodial structure and cell migration, while constitutively active S3A mutants alter migration, adhesion and actin organization [8]. These findings highlight the critical role of phospho-regulation at the leading edge as a key determinant of membrane dynamics, and suggest that it must be tightly controlled. Several kinases have been identified that target cytoskeletal proteins, but much less is known about how phosphatases contribute to their regulation.

Protein phosphatases such as PP1 and PP2A regulate cytoskeletal proteins [9–12], but how they localize to the leading edge, and how their substrate specificity is determined is less clear. Ser/Thr protein phosphatases like PP1 are actually multiple subunit molecular assemblies with both catalytic and regulatory domains, and associated binding proteins. It seems likely that these regulatory partners may influence protein phosphatase localization and substrate binding. One PP1-binding protein in particular, Phosphatase and Actin Regulator 4 (Phactr4), is a good candidate for regulating leading edge protrusion due to its ability to also bind monomeric actin. Previous studies linked Phactr4 deficiency to increased pSer3 cofilin, and compromised lamellipodial structure and neural crest migration [13]. In addition, Phactr4 downregulation and/or dysfunction has been implicated in numerous diseases [14–17]. Thus, Phactr4 may be a critical coordinator of actin dynamics at the leading edge due to its influence on PP1 function. We hypothesized that Phactr4 localizes to branched actin networks and helps coordinate the phases of the protrusion-retraction cycle by regulating the activity of cofilin and other actin-associated proteins at the leading edge.

In the current study, we provide evidence that Phactr4 contributes to lamellipodial maintenance, as Phactr4-deficient macrophages move in an uncoordinated fashion via multiple protrusive fronts. Further investigation of leading-edge dynamics revealed that Phactr4-deficient cells protruded and retracted more dynamically than control cells. These unproductive protrusions result in dysregulated cell migration and inefficient complement-mediated phagocytosis in Phactr4-depleted macrophages. We provide molecular evidence that ezrin hyperactivation is a major element of the Phactr4 deficiency phenotype, indicating that Phactr4 likely coordinates PP1-dependent ezrin inactivation. Finally, we provide novel evidence that Phactr4 interacts with Arp2/3 complex, and that Phactr4 leading edge localization is lost upon Arp2/3 inhibition. These data together suggest that Phactr4 has a significant role in regulating lamellipodial protrusion by coordinating protrusion/retraction cycles via PP1-dependent phospho-regulation. The link between ezrin phosphorylation and Phactr4 may be a relevant component of pathophysiology, as Phactr4 deficiency may correlate with ezrin hyperphosphorylation in human diseases.

## RESULTS

### Phactr4 localizes to macrophage lamellipodia and regulates protrusion number

We started our investigation on Phactr4 in macrophages by determining that it localizes to the cell edge in close proximity to F-actin (**Fig. 1A, quantified in Fig. 1B**), suggesting a role in lamellipodial structure and dynamics. siRNA depletion of Phactr4 in primary macrophages (**Fig. 1C**) led to striking changes to cellular morphology. Macrophages transfected with scrambled siRNA (denoted siCon) presented with a single, broad lamellipodia (**Fig. 1D**). Phactr4 KD (siPh4) macrophages generated numerous branched protrusions and were smaller than their control counterparts (**Fig. 1D, quantified in Fig. 1E**; **Sup. Fig. 1A**).

**Figure 1.**
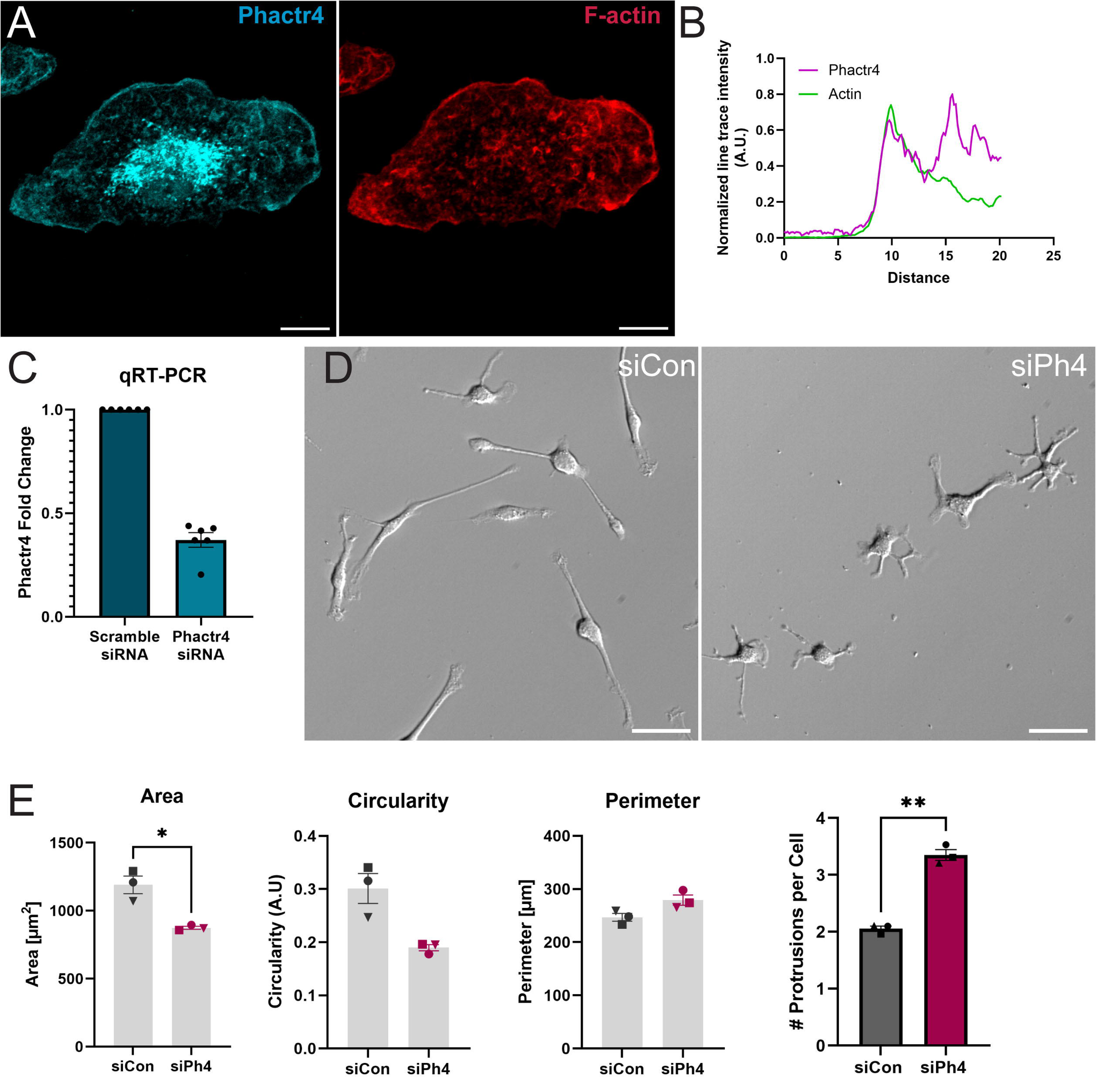
Phactr4 regulates macrophage lamellipodia number and formation. (A) Representative image of a WT macrophage stained for Phactr4 (RRX) and F-actin (Phalloidin 647). Images were acquired using confocal microscopy and are presented as maximum Z projections. Scale bar = 5 µm. (B) Line trace quantification of Phactr4 and F-actin fluorescence intensity along the same line. Fluorescence intensity is normalized to the highest signal along the trace. The line trace demonstrates that Phactr4 and F-actin closely overlap at the cell edge, with Phactr4 peaking slightly ahead of F-actin near the cell membrane. (C) Phactr4 fold change from qRT-PCR. Data represent N = 6 populations of immortalized macrophages, with siPhactr4 expression normalized to the paired siControl population at the time of transfection. (D) Representative relief contrast images comparing macrophages transfected with siControl versus siPhactr4. Cells were plated on 1 µg/µL fibronectin for approximately 10 hours prior to imaging. Scale bar = 50 µm. (E) Quantification of cell area (µm²), circularity, and perimeter (µm) using ImageJ manual selection and measurement functions. Data represent N = 3 independent experiments with 100–130 cells per condition. Statistical analysis was performed using Welsh’s t-test, with *p = 0.0361. (F) Protrusion number was quantified manually, with a protrusion defined as an extension from the primary cell body. Protrusions branching from existing protrusions were not counted. Statistical analysis was performed using Welsh’s t-test, with **p = 0.0014.

### Phactr4 deletion disrupts cell migration and phagocytosis

Since membrane protrusions are altered in siPh4 cells, we hypothesized that protrusion-dependent processes like migration and phagocytosis would also be altered. To analyze cell spreading, we synchronized cellular protrusion by briefly serum starving each population (**Sup. Fig. 1B**). The siPh4 cells start out smaller at baseline, but both populations become smaller during serum deprivation due to retraction of their protrusions (**Sup. Fig. 1B-C**). However, siPh4 cells are unable to spread upon serum re-addition, while siCon cells return to their average baseline size (**Sup. Fig. 1C**). We noted no significant change in perimeter or circularity when comparing within the same conditions (**Fig. 1E, Sup. Fig. 1C**). These data suggest that Phactr4 is fundamentally linked to membrane protrusion during cell spreading.

We next interrogated the role of Phactr4 in cellular behaviors dependent upon membrane protrusion, such as cell migration and phagocytosis. siPh4 macrophages were remarkably less motile than siCon cells in random motility experiments, leading to shorter migration tracks (**Fig. 2A, Sup. Movie 1**). Quantification of these cell tracks revealed that cell speed and distance traveled were both significantly decreased by Phactr4 depletion (**Fig. 2B**). siPh4 cells also moved with less directional persistence (**Fig. 2B**). These data implicate Phactr4 as a fundamental regulator of cell migration. Next, we challenged siCon and siPh4 cells with six-micron pHRodo-iC3b conjugated beads to evaluate how Phactr4 depletion impacts membrane protrusion during phagocytosis. Six-micron beads were chosen to force macrophages to make large, coherent protrusions to internalize phagocytic targets. pHRodo fluorescence is pH-sensitive, and indicates particle loading into the phagolysosome, thereby giving us an unbiased way to image phagocytic completion. As expected, siCon macrophages continuously internalized beads through the 16-hour time course (**Fig. 2C, top;** as indicated by arrow, quantified below**, and Sup. Movie 2**). Conversely, siPh4 macrophages had a significant uptake lag compared to siCon cells, which persisted through the final experimental time point (**Fig. 2C**, top and bottom). In fact, siPh4 macrophages seemed unable to effectively initiate and maintain a phagocytic cup, as we captured numerous examples of beads contacting unresponsive siPh4 macrophages (**Fig. 2C, Sup. Movie 2**). In line with these observations, Phactr4 strongly co-localized with F-actin in iC3b phagocytic cups in control macrophages (**Fig. 2D**). Some siPh4 cells do eventually internalize beads (**Fig. 2C, bottom; Sup. Fig. 2A**). Once internalized, phagolysosomes form and bead fluorescence occurs similarly in both populations (**Sup. Fig. 2B**). These results indicate that the siPh4 defect is specific to bead internalization, rather than impaired intracellular trafficking to the phagolysosome. Together, these data establish that Phactr4 has a strong influence on cellular behaviors that require dynamic membrane protrusion.

**Figure 2.**
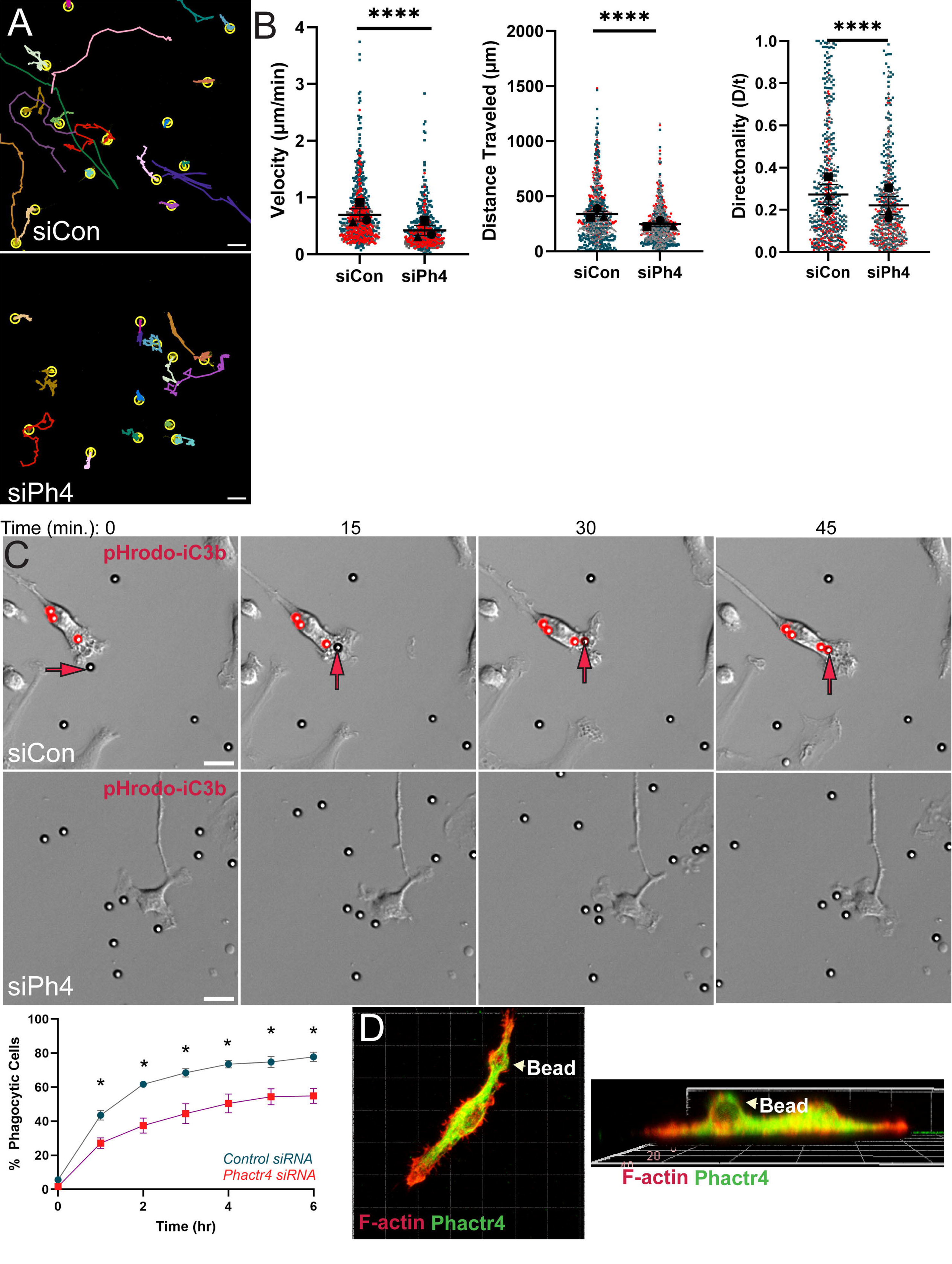
Phactr4 regulates macrophage migration and phagocytosis. (A) Representative tracks of siControl and siPhactr4 immortalized macrophages migrating randomly over a 16-hour period. Tracks were generated using the TrackMate Fiji plugin. Cells were plated on 1 µg/mL fibronectin. Scale bar = 50 µm. (B) Velocity (cell speed, µm/min), persistence (d/T), and total distance traveled (µm) were quantified using TrackMate Fiji. Data are pooled from three independent experiments. Means and standard error of the mean for each experiment are represented with black symbols, all data points are plotted, and each experimental run is color-coded, with the shape corresponding to the mean. Approximately 100 cells per condition per experiment were analyzed, with N = 3 individual experiments. Statistical analysis was performed using Welsh’s t-test, ****p < 0.0001. (C) Representative images of siControl and siPhactr4 immortalized macrophages encountering iC3b-opsonized beads. The leftmost frame shows the first timepoint at 5 hours after bead addition, followed by consecutive images taken every 15 minutes. Images are relief contrast overlaid with the red fluorescence channel. In siControl macrophages, red arrows mark a bead that is ultimately phagocytosed, as indicated by increasing fluorescence intensity over time. In contrast, siPhactr4 macrophages encounter multiple iC3b-opsonized targets within the 45-minute time window but do not internalize any beads in proximity. Scale bar = 20 µm. The percentage of phagocytic cells was quantified by manually counting the number of cells with internalized fluorescent beads per field of view. Each experiment contained 10–15 fields of view, with approximately 100 cells per experiment (N = 3 independent experiments). Phagocytosis plateaued by the 6-hour mark, so quantification was limited to this period. Welsh’s t-test was performed at each hourly time marker from 0 to 6 hours: siCon v siPh44 *p = 0.0183 (1 hr), *p = 0.0306 (2 hr), *p = 0.0391 (3 hr), *p = 0.0380 (4 hr), *p = 0.0285 (5 hr), *p = 0.0171 (6 hr). (D) 3D rendering of a representative WT macrophage after internalizing an iC3b-opsonized bead. Cells were fixed after a 20-minute incubation with pHrodo-labeled beads, and immunofluorescence staining was performed to visualize Phactr4 and F-actin. Images were acquired via confocal microscopy, and 3D rendering was conducted using Avaris software in Zeiss Zen Blue. The left panel shows a bottom-up view of the cell, while the right panel presents a side view, with arrows indicating the position of the 6 µm bead.

### Phactr4 regulates protrusion/retraction dynamics and localizes to retracting membranes

Since Phactr4 depleted macrophages cannot form productive leading-edge protrusions, we hypothesized that altered membrane dynamics would be the root cause of the siPh4 phenotypes. Kymographs reveal that siCon macrophages produce stereotypical large protrusions followed by slow retractions, while siPh4 macrophages constantly protrude in a way that leads to choppy leading-edge dynamics (**Fig. 3A, Sup. Fig. 3A, Sup. Movie 3**). This contrast is further revealed by quantitative measurement of kymographs from each cell. First, the velocity of the siPh4 cell edge is faster during protrusions compared to siCon (**Fig. 3B**), leading to shorter protrusion and retraction durations (**Fig. 3C**), though displacement distance was not affected in a statistically significant fashion (**Sup. Fig. 3B**). These dynamic membrane fluctuations lead to an overall more dynamic cell edge, as evidenced by increased protrusion-retraction frequency in siPh4 macrophages (**Fig. 3D**), and as a result they spend less time in a paused state (i.e. neither protruding or retracting, **Fig. 3E**, examples in **Sup. Fig. 3C**). Next, expression of GFP-tagged Phactr4 was used to gain temporal insight into the role of Phactr4 at the leading edge. Strikingly, GFP-Phactr4 enrichment at the leading edge corresponds strongly with lamellipodial retraction (**Fig. 3F, 3G, Sup. Movie 4**). These GFP-Phactr4 overexpressing cells have a much more dynamic cell edge than GFP-expressing cells as evidenced by enhanced protrusion-retraction cycling (**Fig. 3H**). Time lapse microscopy revealed that GFP-Phactr4 localization to the cell edge dynamically altered lamellipodial structure, and that at times GFP-Phactr4 signal spread laterally across the cell edge as it became remodeled (**Fig. 3I, red arrows**). The striking phenotypes from disruption (siPh4) and overexpression (GFP-Ph4) suggest that Phactr4 activity must be tightly controlled, and is likely regulated in a context-dependent fashion. These data also suggest that Phactr4 regulates leading edge actin-associated proteins that determine lamellipodial structure and dynamics.

**Figure 3.**
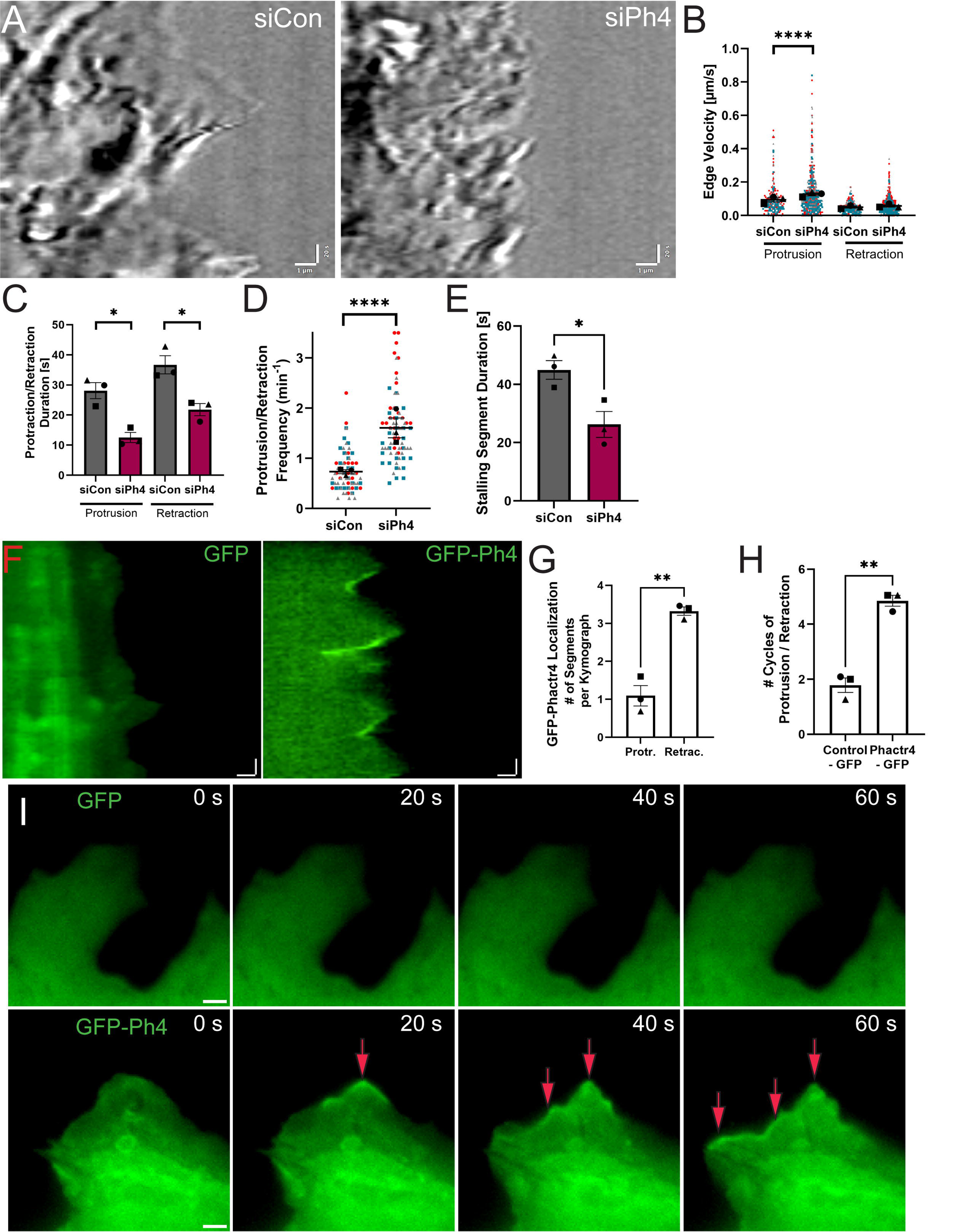
Phactr4 controls macrophage protrusion/retraction dynamics. (A) Example kymographs generated using CellSens software. Scale bars: horizontal = 1 µm, vertical = 20 seconds. (B) Kymography analysis using CellSens quantified velocity for each segment of the kymograph. The corresponding DIC video and kymograph were used to map each segment to either the protruding or retracting phase of membrane movement. Individual velocity values for protrusive and retracting segments are plotted. Statistical analysis was performed using Welsh’s t-test, ****p < 0.0001. (C) Duration (in seconds) of individual kymograph segments was quantified using CellSens kymography analysis. Means and standard error of the mean are represented with black symbols. Statistical analysis was performed using Welsh’s t-test, with *p = 0.0115 for protrusion duration and *p = 0.0198 for retraction duration. Data represent N = 3 independent experiments, with 30–35 kymographs analyzed per experiment per condition. (D) Protrusion/retraction cycles were defined by identifying peaks and troughs in the kymograph. The number of cycles during the 5-minute video was manually counted and divided by 5 to determine the frequency of cycles per minute. Data represent N = 3 independent experiments, with 30–35 kymographs analyzed per condition per experiment. Means and standard error of the mean for each experiment are represented with black symbols, all data points are plotted, and each experimental run is color-coded, with the shape corresponding to the mean. Statistical analysis was performed using Welsh’s t-test, ****p < 0.0001. (E) Some kymographs contained segments where the cell edge exhibited ambiguous behavior, making it difficult to categorize movements as protrusive or retractive. These segments were labeled as “stalling segments,” and the total stalling time per kymograph was quantified. Means and standard error of the mean for each experiment are represented with black symbols. Statistical analysis was performed using Kruskal-Wallis with Dunn’s multiple comparisons test. *p = 0.0263. (F) Example kymographs generated using CellSens software kymography tool. Scale bars: horizontal = 1 µm, vertical = 20 seconds. (G) Quantification of protrusion-retraction cycle frequency per kymograph. Kymographs were selected from the most dynamic regions of fibroblasts. A single protrusion-retraction cycle was defined by the peak and trough of the cell edge, visualized using diffuse GFP signal and corresponding DIC images. The number of cycles was manually counted to calculate frequency per minute. Data represent N = 3 independent experiments, with 10–15 kymographs analyzed per condition per experiment. Each experiment is represented by a different shape, with mean and standard error of the mean indicated. Statistical analysis was performed using Welsh’s t-test, **p = 0.0001. (H) GFP-Phactr4 localization was analyzed by defining each kymograph segment as either protrusive or retracting, and binary localization was determined based on whether GFP-Phactr4 was present in one of these phases. Data were quantified per kymograph, **p = 0.0070, Welsh’s t-test. (I) Time series of an example protrusion in GFP-control (top) and GFP-Phactr4 (bottom) fibroblasts. Frames are shown at 20-second intervals. GFP-Phactr4 exhibits dynamic relocalization during membrane movement, while GFP-control remains diffuse in the cytoplasm with no observable changes over the time series. Red arrows indicate laterally-spreading GFP-Phactr4 signal across the cell edge.

### PP1 localization and ezrin phosphorylation are altered in siPh4 macrophages

We turned our attention to PP1 as it is a known Phactr4 binding protein that dephosphorylates actin-associated proteins. A subset of PP1 localizes to Arp2/3 complex-containing siCon membrane protrusions, but appears reduced in siPh4 macrophage protrusions (**Fig. 4A**). We hypothesized that PP1 activity would be dysregulated in siPh4 cells, causing hyperphosphorylation of several cytoskeletal regulators relative to siCon. While these experiments failed to detect a consistent trend for p-MLC, p-cofilin and p-Arp2, all three trended upward in the siPh4 cells (**Sup. Fig. 4A**). Total vinculin levels trended slightly lower (**Sup. Fig. 4A**). In contrast, increased ezrin phosphorylation stood out as a robust and reproducible change in the siPh4 cells (**Fig. 4B**). Adding support for this regulatory connection, ezrin and Phactr4 were also found in close proximity to each other via proximity ligation assay (**Sup. Fig. 4B, 4C**). Their proximity was not affected by Arp2/3 inhibition (**Sup. Fig. 4B, 4C**), suggesting that lamellipodial localization is not required for this function of Phactr4. Ezrin is a key member of the <u>E</u>zrin <u>R</u>adixin <u>M</u>oesin (ERM) family [18, 19], which together serve as critical linkers of the actin cytoskeleton and the plasma membrane. Its activation via phosphorylation at the T567 residue promotes actin-membrane interactions [20–23], while dephosphorylation is required for membrane detachment preceding Arp2/3-dependent protrusion [24]. Ezrin hyperactivation in the siPh4 cells could restrict protrusion dynamics, thereby giving rise to the phenotypes seen in the siPh4 cells. To test whether inhibiting ezrin activity could rescue the siPh4 membrane protrusion defect, we treated siPh4 macrophages with an inhibitor that prevents T567 phosphorylation of ezrin. Inhibitor-treated siPh4 cells regained cell edge dynamics reminiscent of siCon cells (**Fig. 4D, Sup. Movie 5**). Indeed, quantification revealed that inhibitor-treated siPh4 cells regained protrusion-retraction dynamics (**Fig. 4E**, top), displacement (**Fig. 4E**, bottom), and duration (**Sup. Fig. 4C**) similar to siCon. As a whole, these data support the notion that Phactr4 delivers PP1 to the leading edge, which supports productive membrane protrusion in part by influencing the phospho-regulation of ERM proteins. Next, we wanted to understand how Phactr4 is recruited to the cell edge.

**Figure 4.**
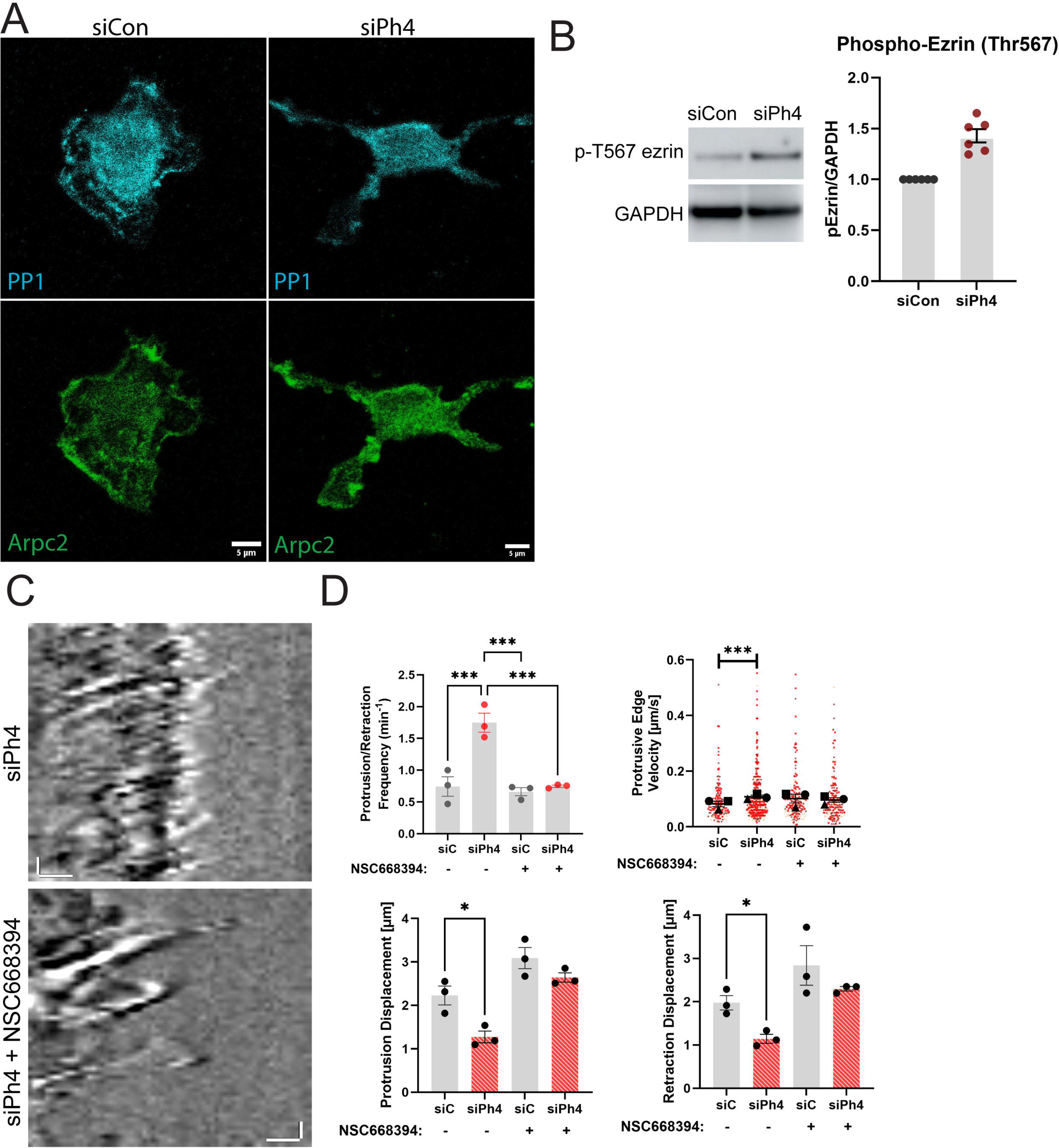
Phactr4-mediated Ezrin dephosphorylation supports normal protrusion formation. (A) Representative images of siControl and siPhactr4 macrophages stained for PP1 and Arpc2. Scale bar = 5 µm. (B) Left: Representative western blot images showing phosphorylated Ezrin (p-Ezrin, Thr567) levels in siControl and siPhactr4 conditions. GAPDH was used as a loading control. Representative blot from N = 6 independent experiments. Right: Quantification of p-Ezrin signal after normalization to GAPDH. Means and standard error of the mean are represented. (C) Example kymograph of siPhactr4 macrophages treated with either DMSO control or inhibition of Ezrin Thr567 phosphorylation using 100µM NSC668394. Scale bars: horizontal = 1 µm, vertical = 20 seconds. (D) Quantification of dynamic edge parameters from kymography analysis, including protrusion-retraction cycle frequency, protrusion and retraction duration, and displacement. Kymographs were chosen from the most dynamic regions of fibroblasts, with protrusion-retraction cycles defined as the peak and trough of the cell edge, visualized using corresponding DIC images. The number of cycles was manually counted to calculate frequency per minute. Data represent N = 3 independent experiments, with 10–15 kymographs analyzed per condition per experiment. Means and standard error of the mean for each experiment are represented. For protrusive edge velocity analysis, means are represented with black symbols, all data points are plotted, and each experimental run is color-coded, with the shape corresponding to the mean. Statistical analysis was performed using one-way ANOVA, ***p = 0.0005. Protrusive edge velocity was analyzed using Kruskal-Wallis with Dunn’s multiple comparisons test. siControl vs. siPhactr4 ***p = 0.0004, which was lost after Ezrin inhibition (siControl NT vs. siPhactr4 + NSC668394, ns = 0.6743). Protrusion displacement *p = 0.0275, retraction displacement *p = 0.0191.

### Phactr4 is recruited to the leading edge via interaction with the Arp2/3 complex

It is readily apparent that Phactr4 and the Arp2/3 complex co-localize at the leading edge of the cell (**Fig. 5A**). Line traces reveal that the leading-edge peak for Phactr4 is almost perfectly overlaid with the leading-edge peak for Arp3 (**Fig. 5B**), suggesting a tight coordination in their localization. From these data we hypothesized that Arp2/3 recruits Phactr4 to the leading edge. Consistent with our hypothesis, treating GFP-Phactr4 expressing cells with the small molecule Arp2/3 inhibitor CK-666 delocalized GFP signal from the cell edge (**Fig. 5C; Sup. Movie 6**). We used CK-666 washout experiments to further test this idea using Phactr4 antibody staining as a readout. Notably, washing out the CK-666 leads to both recovery of leading edge lamellipodial structure and enhanced Phactr4 localization to the leading edge during the coordinated re-spreading of the cell after CK-666 (**Sup. Fig. 5A, Sup. Fig. 5B**). Both PP1 and Arp2/3 co-IP with Phactr4 (**Fig. 5D**), in line with the hypothesis that a physical interaction with Arp2/3 localizes Phactr4 to the edge. Finally, proximal ligation assays demonstrate that the Arp2/3-Phactr4 interaction occurs at the leading edge and throughout the cell body when Arp2/3 is active (**Fig. 5E, 5F**, no treatment and CK-666 washout conditions). Just as Phactr4 is delocalized during acute CK-666 treatment, the interaction between Arp2/3 and Phactr4 is disrupted upon Arp2/3 inhibition (**Fig. 5F, 5G**). This body of work identifies Phactr4 as a novel regulator of cell edge dynamics that physically interacts with active Arp2/3 complex and ezrin, and modulates the function of actin-associated proteins like ezrin via phosphoregulation.

**Figure 5.**
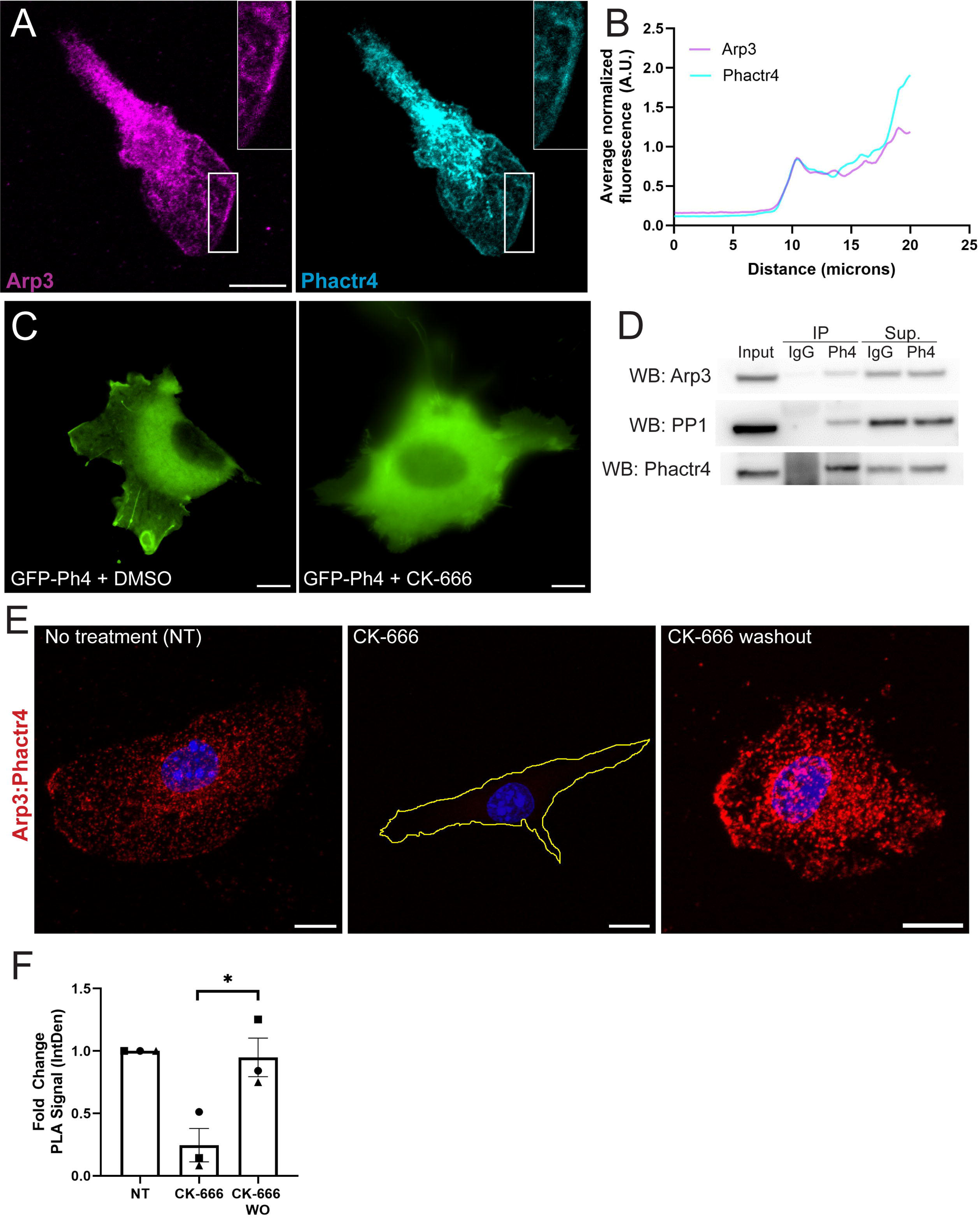
Phactr4 is recruited to the cell’s leading edge by active Arp2/3 complex. (A) Representative image of a WT macrophage stained for Arp3 and Phactr4. (B) Line trace analysis of fluorescence intensity along both channels, taken along a line intersecting the lamellipodia of a macrophage. Fluorescence intensity is normalized to the maximum signal at the cell membrane. (C) Representative images of GFP-Phactr4-expressing macrophages treated with media containing either 100 µM CK-666 (right) or DMSO control (left). Cells were plated on 10 µg/mL fibronectin and transfected two days prior to imaging. Treatment was conducted for 1 hour before imaging. Scale bar = 10 µm. (D) Western blot images from co-immunoprecipitation (co-IP) assay showing bands for Arp3, PP1, and Phactr4. The left lane is the loading control. IP lanes indicate samples precipitated using either IgG control or Phactr4 antibody. The right two lanes show supernatant from IgG and Phactr4 IP samples. (E) Representative confocal images of proximity ligation assay (PLA) signal from the Duolink PLA assay, shown as maximum projections from a z-stack. Fluorescent puncta represent individual localizations of Arp3-Phactr4 interactions. Cells were either untreated (left), treated with 100 µM CK-666 for 2 hours (middle), or treated with 100 µM CK-666 for 2 hours followed by a 10-minute media washout (right) before fixation and PLA assay. Scale bar = 10 µm. (F) Quantification of PLA fold change. The relief contrast channel was used to outline the cell perimeter, and this selection was mirrored onto the fluorescent channel in ImageJ. The integrated density (InDen) of each cell was measured. Each experiment included ∼15 cells, with N = 3 independent experiments. PLA signal was normalized to the no-treatment control. Means and standard error of the mean are represented. Statistical analysis was performed using Welsh’s t-test between CK-666 vs. CK-666 washout, *p = 0.0272.

## DISCUSSION

Our findings highlight Phactr4 as a critical regulator of lamellipodial protrusions and macrophage function. Phactr4’s activity-dependent interaction with the Arp2/3 complex likely coordinates PP1 activity with the branched actin network to ensure that phospho-regulation at the leading edge is properly orchestrated. Phactr4 also interacts with ezrin, and Phacr4 deficiency correlates with ezrin hyperphosphorylation. Ezrin hyperphosphorylation also likely plays a key role in the lamellipodial dysfunction seen in the Phactr4 deficient cells, as this phenotype is corrected upon Ezrin inhibition. Each of these findings have potentially major impacts on our understanding of lamellipodial structure and dynamics and the interplay between ERM proteins and Arp2/3 complex, as well as presenting new avenues for better understanding human disease.

Our findings provide evidence that Phactr4 contributes to the formation of stable, functional lamellipodial protrusions. The present study and previous literature agree that Phactr4 deficiency leads to multiple, small lamellipodial protrusions that lead to uncoordinated migration [13]. One fundamental reason for this could be increased membrane coupling to the actin cortex through Ezrin/Radixin/Moesin (ERM) proteins, which could render the siPh4 cell membrane more resistant to large deformations. Compelling recent work actually demonstrates that in order for an organized lamellipodial protrusion to form, Ezrin release from the membrane precedes Arp2/3-dependent protrusion [24]. In this sense, Phactr4 may be acting ‘pre-protrusion’ to define the site of a future lamellipodia by locally relaxing membrane tension via ERM inhibition. Phactr4-deficient cells may only be able to initiate disorganized lamellipodia at random patches of low-tension membrane. Increased integrin activation in siPhactr4 cells [13] could also contribute to lamellipodial dysfunction, possibly by increasing contractility and/or further resisting membrane deformation [25]. In addition, Phactr4 may help position PP1 near other phospho-regulated proteins that affect lamellipodial initiation and architecture, such as Lamellipodin [26, 27], Ena/VASP [28, 29] and SCAR/WAVE [30, 31]. Phactr4 could activate or restrict these factors spatially or temporally to help balance the protrusion-retraction cycle that drives lamellipodia forward.

GFP-Phactr4’s enrichment in membrane retractions suggests that Phactr4 plays an active role in coordinating lamellipodial dynamics. This dovetails nicely with Phactr4’s established function in cofilin activation via dephosphorylation, which would be expected to locally stimulate actin severing [10, 32, 33]. Our bulk western blot analyses could not definitively confirm that cofilin phosphorylation was increased in Phactr4-deficient macrophages. However, it certainly could be the case that the spatiotemporal activation of cofilin does not occur efficiently in these cells, which could be missed by population-level experiments like western blots. We also noted that the Phactr4-deficient cells cycled faster between protrusion and retraction, and demonstrated a characteristic ‘choppy’ cell edge compared to siCon. The siPh4 protrusion phenotype was corrected by Ezrin inhibition, which suggests that the disordered dynamics at the cell edge could be due in part to altered membrane and/or cortical tension. Thus, there are likely multiple layers of Phactr4 function yet to be uncovered.

We were surprised to find that active Arp2/3 complex recruits Phactr4 to the cell edge. Given Phactr4’s enrichment in membrane retractions, our initial suspicion was that Phactr4 would directly impact Arp2/3 complex function. This led us to interrogate whether Phactr4 deficiency correlated with an increase in Arp2 phosphorylation, potentially enhancing the nucleation rate of branched actin in the siPh4 cells [34, 35]. As with p-cofilin, we were unable to see a consistent trend with p-Arp2 in our siPh4 cells at the bulk protein level. However, local p-Arp2 levels could be higher at the leading edge lamellipodia in siPh4 cells compared to siCon. The impaired protrusion phenotype of the siPh4 cells may argue against enhanced Arp2/3 complex nucleation. Beyond Arp2/3 complex itself, the stability of the branched actin network is determined by a variety of factors that positively or negatively regulate the branched actin network. Many negative regulators of Arp2/3 complex have been identified, including coronins [36, 37], GMF [38], and Arpin [39], among many others. It is possible that Phactr4’s localization to Arp2/3 complex influences these negative regulators rather than Arp2/3 itself, perhaps via altered phospho-regulation. Coronin 1B S2E mutation disrupts its interaction with Arp2/3 complex, leading to impaired motility, while S2A mutants demonstrate increased interaction with Arp2/3 which leads to enhanced motility and membrane ruffling [40]. In a similar fashion, GFP-Phactr4 expression in fibroblasts in the present work led them to become much more protrusive, with structures appearing similar to membrane ruffling, suggesting that Phactr4 similarly enhances actin dynamics and membrane ruffling at the cell edge. Active Arp2/3 complex may serve as a scaffold to position Phactr4-dependent delivery of PP1 to suppress these phosphorylation events and thereby facilitate branched actin turnover. As important as Phactr4 seems to be for lamellipodial dynamics, we also provide evidence that there are Arp2/3 complex-independent roles for Phactr4 in the cell. Phactr4 and Ezrin remain in close proximity during CK-666 treatment, indicating that Ph4 may maintain a balance of Arp2/3-dependent and – independent functions that can be shifted in a context-dependent fashion.

At the cellular level, Phactr4 deficiency leads to defects in phagocytosis and whole cell motility which both require dynamic, coordinated membrane protrusions. siPh4 cells fail to orient polarized protrusions and migrate persistently on fibronectin. The siPh4 cell leading edge often collapses prematurely, forcing the cell to repeatedly reorient in an uncoordinated fashion. This pattern suggests that Phactr4 is essential not only for protrusion stability but also for establishing and/or maintaining front-rear polarity, a prerequisite for efficient directional migration. Phactr4 mutant fibroblasts fail to reorient the microtubule-organizing center (MTOC), which is a well-established marker of front-rear polarity [13]. As suggested previously [41], the Phactr4-PP1 interaction is also potentially relevant in this context, as PP1 has been identified as a Par complex regulator [42, 43]. The Par complex coordinates cytoskeletal remodeling, integrin trafficking, and centrosome reorientation [44] at the leading edge. Loss of PP1 activity destabilizes the Par complex, impairing its ability to polarize cells during migration [45]. It will be interesting to assess whether disrupted Par complex function plays a role in the siPh4 macrophage phenotypes. Inefficient protrusion and increased F-actin coupling to the membrane via Ezrin activation are also highly relevant to the siPh4 phagocytic defect. siPh4 macrophages interact with beads, but fail to internalize them even after prolonged interaction. Transient extensions in siPh4 macrophages may lack sufficient protrusive force, retract prematurely, fail to properly regulate integrin signaling, or exhibit enhanced mechanical tension that limits membrane deformation.

The current study may also shed light on why Phactr4 has been linked to numerous human diseases. In particular, the link between Ezrin and Phactr4 may be relevant to tumor pathogenesis. Phactr4 deletions and ezrin activation have been noted in several cancers, including breast, colorectal, lung, ovarian, and renal tumors [15, 46–49]. It is possible that selective downregulation of Phactr4 in monocyte-derived tumor-associated macrophages (TAMs) could lead to migratory defects that prevent them from effectively penetrating the tumor microenvironment, as well as to phagocytic defects that prevent tumor cell clearance. Upon their recruitment to tumors, newly activated macrophages are reprogrammed by tumor-derived factors, including TGF-β, IL-10, and hypoxic signals, into a phenotype that aligns more closely with tissue repair and immune suppression than robust tumor clearance [50]. Decreased TAM-mediated phagocytic clearance of complement-labeled tumor cells is a feature of this process [51], once more echoing the siPh4 phenotype. Phactr4 may also fundamentally influence immune activation. A recent study reported that Phactr4 upregulation in the hippocampus in a chronic stress-induced depression model correlates with NF-kB induction and IL-1β secretion [52]. Phactr4 downregulation in this context decreased neuroinflammation, improved synaptic plasticity, and moderated depression-like behaviors in rats [52]. It will be interesting to determine whether the link between Phactr4 and inflammation is also linked to cytoskeletal disruption. In summary, our work establishes Phactr4 as a critical player linking actin cytoskeleton dynamics to immune regulation. In shedding light on this understudied regulator, we aim to stimulate future systematic investigation of phosphoregulatory immune networks in pathological settings such as the tumor microenvironment and neurological disorders.

## MATERIALS AND METHODS

*Cell Culture and Treatments*. <u>Murine bone marrow derived macrophages</u>: Bone marrow-derived macrophages (BMDMs) were isolated from the femurs of wild-type C57BL/6 male mice aged 4– 6 months. Briefly, femurs were excised and both ends were trimmed using a razor blade. The bone marrow was then flushed out with PBS supplemented with 2% fetal bovine serum using a 22G needle attached to a 10mL syringe inserted into the femur. The resulting cell suspension was passed through a 70-µm filter (STEMCELL Cat. #27260) to enrich for hematopoietic progenitors. The filtered cells were centrifuged at 1000 rpm for 5 minutes, and the pellet was resuspended in Macrophage Media (MM) consisting of DMEM (4.5 g/L D-Glucose, L-Glutamate, Sodium Pyruvate, minus phenol red; Corning Cat. #31053036), 10% FBS (Sigma-Aldrich Cat. #12306C), 1% Glutamax (ThermoFisher Scientific Cat. #35050061), and 30% (v/v) L949 cell-conditioned media. Cells were then cultured in 10-cm dishes (CellTreat Cat #22962) at 37°C with 5% CO₂ and 90% humidity. BMDMs were maintained in MM and passaged for experimental use upon reaching 60–90% confluency. To passage cells, they were first washed with 1X PBS and then incubated for 10 minutes at 4°C in prechilled 0.5 mM ethylenediaminetetraacetic acid (EDTA, Invitrogen Cat. #AM9260G). After removing the EDTA, BMDMs were gently scraped into 1–3 mL of macrophage media (volume adjusted based on desired confluency) and subsequently replated for further experimentation. <u>Immortalized murine bone marrow derived macrophages</u>: Immortalized macrophages were derived from a mixed strain of mice (predominantly C57BL/6) carrying a conditional Arpc2 allele, in which LoxP sites flank exon 8 of the gene encoding the p34/Arpc2 subunit of the Arp2/3 complex [53]. Additionally, these cells harbor Ink4a^−/− and Arf^−/− alleles, a defined genetic modification that prevents senescence and supports sustained proliferation, rather than arising from spontaneous oncogenic events. They also express a Rosa26-CreER transgene, enabling conditional deletion of Arpc2 exon 8 in the presence of tamoxifen. Since no tamoxifen was administered in our experiments, the cells were considered functionally wild-type throughout their culture. This specific combination of genetic alterations has been previously described [53]. In addition, an alternative immortalized macrophage cell line, RAW 264.7 (ATCC Cat. TIB-71), was maintained under the same culture conditions as the primary macrophage types. <u>Dermal fibroblast culture line</u>: JR20 cells are a stable clonal fibroblast line derived from perinatal mouse dermis [53]. These cells also are Ink4a/Arf knockout, and harbor the floxed Arpc2 allele. JR20 cells were cultured in DMEM (4.5 g/L D-Glucose, L-Glutamate, Sodium Pyruvate, minus phenol red; ThermoFisher Cat. #31053028), supplemented with 10% FBS (Sigma-Aldrich #12306C) and 1% Glutamax (ThermoFisher Scientific Cat #35050061). Cells were passaged upon reaching 60-90% confluency using Trypsin-EDTA (Caisson Labs Cat. #TRL02) incubation at 37 °C

*Fibroblast transfection*. For each transfection of GFP-tagged protein with Lipofectamine 3000 (Thermo, L3000001), 1 μg of plasmid DNA was combined with 2 μL of P3000 reagent in 100 uL Opti-MEM reduced serum medium (ThermoFisher Cat. #31985062). Separately, 2 μL of Lipofectamine™ 3000 was diluted in 100uL Opti-MEM. These two solutions were mixed and incubated at room temperature for 15 minutes to allow the formation of DNA–lipid complexes. The transfection mixture was then added dropwise to fibroblasts seeded at approximately 70– 80% confluency in 35-mm dishes (STEMCELL Cat. #27150). The GFP-Phactr4 plasmid [54] was kind gift from Dr. Maria Vartiainen, University of Helsinki. The GFP vector control was cloned and then re-inserted into the same plasmid backbone, which served as a negative control.

*Inhibitors and other reagents*. CK-666 (Sigma Aldrich, 182515); NSC668394 (MedChemExpress, HY-115492); CellBrite Orange (Biotium, 30022); Alexa Fluor 647 Phalloidin (Invitrogen, A22287).

*Western blotting*. <u>Cell lysis</u>: Cells were plated at a density of 200,000–300,000 per well in 3.5 cm plastic culture dishes (StemCell Tech, Cat. 27150) and allowed to spread overnight before starting treatments. After treatments, cells were washed three times with ice-cold PBS and lysed on ice using RIPA buffer (Sigma, Cat. R0278-50ML) supplemented with 1X protease inhibitor (Thermo Fisher Scientific A32953) and 1X phosphatase inhibitor (Sigma, Cat. 4906845001). Cells were scraped using a cell lifter (CellTreat, Cat #229305) and collected in microcentrifuge tubes, which were incubated on ice for 15 min. Lysates were centrifuged at 15,000× g for 15 min at 4°C, and the supernatant was collected while the pellet was discarded. Protein concentration was measured using Precision Red (Cytoskeleton, Cat. ADV02-A) with BSA standards prepared in RIPA buffer to generate a standard curve for quantification via plate reader. <u>SDS-PAGE</u>: For electrophoresis, protein samples (8–15 µg) were mixed with 1X Bolt LDS Sample Buffer (ThermoFisher Scientific B0007) and 1X NuPAGE Sample Reducing Agent (Novex NP0009), with remaining volume adjusted using excess RIPA buffer to ensure equal protein concentration across samples. Samples were briefly vortexed, heated to 90°C for 8 min, and centrifuged at max speed for 1 min before loading onto a Bolt 4–12% Bis-Tris protein gel (Thermo Fisher Scientific NW04125BOX). Electrophoresis was performed in 1X NuPAGE MES SDS Running Buffer (Thermo Fisher Scientific NP0002) at 200 V until the dye front reached the bottom of the gel. For protein transfer, a traditional wet transfer onto a 0.45 µm PVDF membrane (Bio-Rad 1620174) was performed on ice for 90 min at 0.5 A using prechilled 1X transfer buffer (19.70 g Trizma HCl, 15.14 g Trizma base, 142.63 g glycine per liter, adjusted to pH 8.3, supplemented with methanol to a final concentration of 20%). <u>Transfer and Imaging</u>: Protein transfer for co-immunoprecipitation samples utilized Power Blotter semi-dry transfer system (Invitrogen PB0010) using the mixed-range molecular weight transfer program (1.3 amps for 10 min). Membranes were blocked in 5% non-fat dry milk in TBST (1X Tris-buffered saline + 0.1% Tween-20) for 1 h at room temperature, followed by incubation with primary antibodies diluted in either 1% BSA or 2% milk in PBST or TBST overnight at 4°C with gentle agitation. Membranes were washed three times for 5 min each with PBST and incubated for 30 minutes to 2 hours at room temperature with HRP-conjugated secondary antibodies (1:10,000 dilution in 5% milk in TBST). After three additional 5-min washes in PBST, HRP signal was detected using SuperSignal West Pico Chemiluminescent Substrate (Thermo Fisher Cat. #34580) and imaged using GE Amersham 680 Imager. <u>Quantification</u>: For quantification, band intensities were analyzed using FIJI/ImageJ software, ensuring that identically sized regions were used for each protein of interest. All values were normalized to GAPDH as a loading control. Uncropped blots can be found in Supplemental Figures.

*Indirect Immunofluorescence staining*. Sterile 12 mm round coverslips (Electron Microscopy Sciences, Cat. 72291-02) were coated with 400uL of Fibronectin (10 µg/mL) (ThermoFisher Scientific, Cat. 33016015) for 1 hour at 37 °C and then washed with 3 consecutive washes of sterile 1x PBS. Cells were plated at a density of 5,000–10,000 cells per coverslip in a 24-well plate and allowed to adhere at 37 °C, 5% CO₂, and 90% humidity for 4 hours to overnight, depending on the experiment. Cells with no treatment were fixed the same day as plating and cells for experiments utilizing chemical inhibitors were treated and fixed the following day. After treatment with Ezrin Inhibitor (NSC668394 MedChemExpress Cat. HY-115492) or Arp2/3 Inhibitor (CK-666 Abcam Cat. ab141231), cells were washed with room-temperature PBS and fixed with cold 4% paraformaldehyde (PFA) in PBS for 10 minutes at room temperature under a fume hood. Cells were then washed three times with PBS before being permeabilized with 0.1% Triton X-100 (ThermoFisher Scientific, Cat. A16046) in PBS for 5 minutes at room temperature. After three additional PBS washes, cells had a 30 minute incubation in a blocking solution containing 5% Bovine Serum Albumin (BSA, Jackson Immunoresearch Cat. 001-000-161) and 5% Normal Goat Serum (NGS, Jackson Immunoresearch Cat. 115-035-068). Primary antibodies (used at 1:100-1:200) were diluted in 1% BSA and incubated with the cells for 1 hour at room temperature. After incubation, cells were washed three times with PBS and incubated with fluorescent secondary antibodies (1:500 in 1% BSA) and Alexa Fluor™ 647 (ThermoFisher Scientific Cat. A22287) (1:500 in 1% BSA) for 30 minutes at room temperature. Cells were then stained with Hoechst (ThermoFisher Scientific Cat. #62249) diluted 1:20,000 in PBS for 5 minutes, followed by one additional PBS wash. Coverslips were mounted onto microscopy slides using 4 µL Fluoromount G (Electron Microscopy Sciences Cat. #17984-25) and allowed to set for 10 minutes in the dark before sealing with nail polish.

*Duolink PLA + Analysis*. The Duolink® Proximity Ligation Assay (PLA) (Sigma-Aldrich Cat. # DUO92101) was performed according to the manufacturer’s instructions. 10,000 cells were plated on fibronectin-coated coverslips in 24-well cell culture plates and treated with 500 µM CK-666, with or without subsequent washout. Following treatment, cells were fixed with 4% paraformaldehyde (PFA) for 15 minutes at room temperature. Cells were then incubated with primary antibodies, anti-Phactr4 (rabbit, 1:500) and anti-Arp3 (mouse, 1:500), for 15 minutes at 37°C followed by an additional 45 minutes at room temperature. After primary antibody incubation, PLA probes (PLUS and MINUS) were applied according to the Duolink protocol, followed by ligation and amplification steps to generate the fluorescent signal at sites of protein-protein interaction. Slides were imaged using a Zeiss 700 confocal microscope with a ×63 oil objective. Z-stacking was performed, with each optical section consisting of ten to fifteen 0.39-μm-thick slices. For quantitative analysis, the mean fluorescence intensity of the PLA signal was measured within the outlined cell area. Relief contrast images of the cells were acquired and used to delineate cell boundaries in ImageJ, ensuring accurate measurement of fluorescence intensity within individual cells. Technical negative controls were performed, including conditions with single primary antibodies and PLA probes only, confirming no detectable background signal.

*Microscopy (fixed samples) and analysis*. <u>Epifluorescence</u> imaging of fixed samples was performed at room temperature using an Olympus IX83 microscope equipped with an X-cite 120 LED Boost light source (Excelitas Technologies) for fluorescence illumination. Images were acquired with a Hammamatsu digital camera (C13440-20CU), and, depending on experimental requirements, 20x, 40x, or 100x objectives were utilized. <u>Confocal</u> imaging was performed using a Zeiss LSM 700 confocal microscope at the USUHS Biological Instrumentation Center (BIC) to obtain high-resolution images of fixed samples with a 63x oil immersion objective. <u>Cell morphology analysis</u>: Images acquired on an Olympus microscope were imported into ImageJ for analysis. Depending on the experiment, either the F-actin channel or relief contrast imaging was used to visualize cell boundaries. The free-hand selection tool was employed to manually trace the exact perimeter of each cell. Measurements, including area, perimeter, and circularity, were then calculated using ImageJ’s built-in measurement functions. <u>Phactr4 cell edge analysis</u>: Images were imported into ImageJ for analysis of Phactr4 protein localization along the cell edge. In macrophages, Phactr4 is typically localized in continuous segments, with some cells exhibiting multiple segments of enrichment. Using the free-hand selection tool on the DsRed channel, individual segments of Phactr4 localization along the cell perimeter were manually traced for each cell. Separately, the entire cell boundary was outlined using the F-actin staining with the same tool to determine the total cell perimeter. The cumulative lengths of all Phactr4-positive segments were then divided by the total cell perimeter, yielding the percentage of the cell edge exhibiting Phactr4 localization. <u>Protein co-localization via line trace:</u> A 20-micron long line was drawn in FIJI such that its midpoint intersected perpendicularly with the front edge of the cell. The fluorescence intensity across this region of interest was quantified in FIJI for all fluorescent channels. The peak leading-edge fluorescence was defined as the highest fluorescent peak value closest to the cell edge. This was normalized to 1 for each individual line trace. The normalized line traces for each channel were then averaged, and the result was plotted in Graph Pad Prism.

*Live cell imaging and analysis*. <u>Random migration assay</u>: Live migration assay was conducted using glass-bottom four-chamber dishes (Cellvis, Cat. C4-1.5H-N) that were pre-coated with 1 µg/mL fibronectin in sterile PBS and incubated at 37°C for 1 hour. Following incubation, the coating solution was aspirated, and the surfaces were rinsed three times with sterile 1X PBS before adding fresh macrophage media to each chamber. The plates were then kept in the incubator until cell seeding. Macrophages were plated at a density of 5,000 cells per well and allowed to adhere for 4 hours before labeling. To facilitate visualization, cells were incubated with a 1:1000 dilution of CellBrite Cytoplasmic Membrane Dye Orange (Biotium #30022) in macrophage media for 1 hour at 37°C. Following the labeling process, the media was replaced with fresh macrophage media, and the dish was transferred to a Tokai Hit stage-top environmental chamber set to 5% CO₂ and 37°C for live imaging. Imaging was performed using an IX83 Olympus microscope with a 20x objective, capturing relief contrast and DsRed fluorescence. A total of 10–15 positions per chamber were imaged every 10 minutes for at least 16 hours. <u>Migration analysis</u>, including measurements of migration velocity, displacement, and persistence, was conducted in FIJI as previously described [55]. Track information was generated using the TrackMate plugin and subsequently processed with the Chemotaxis plugin to obtain individual track values for distance traveled, persistence, and velocity. <u>Phagocytosis assay</u>: Glass-bottom four-chamber dishes (Cellvis, Cat. C4-1.5H-N) were coated with 10 µg/mL fibronectin in sterile PBS and incubated at 37°C for 1 hour. The ECM solution was aspirated, and the surfaces were rinsed three times with sterile 1X PBS. Fresh macrophage media was then added to each chamber, and the plates were kept in the incubator until cell seeding. A total of 5,000 macrophages were seeded per chamber and allowed to adhere for 4 hours before being placed in a Tokai Hit stage-top environmental chamber set to 5% CO₂ and 37°C for live imaging. pHrodo-labeled 6 µm beads were conjugated with iC3b using the pHrodo iFL Red Microscale Protein Labeling Kit (Thermo Fisher, Cat. P36014), following the manufacturer’s protocol. Unpolarized macrophages were treated with 5 µL of the ic3b-bead mixture per chamber and immediately imaged using an IX83 Olympus microscope in the environmental chamber. Live-cell imaging was performed using a 20X objective over a period of 16 hours, with images acquired every 15 minutes. For each condition, 10–15 fields of view were captured per well. Cells were imaged using relief contrast to visualize morphology and the DsRed channel to detect phagocytic activity. <u>Phagocytosis analysis</u> was performed using FIJI. The number of cells per field of view was manually counted using the relief contrast channel, and cells positive for phagocytosis were identified based on the presence of DsRed fluorescence within the cell boundary. Phagocytosis plateaued at 6 hours, so quantification was limited to this time point. Additionally, Olympus CellSens software was used to perform automated quantification of individual phagosome size.

*Kymography and analysis*. Cell edge tracking and kymography analysis were performed using live imaging to assess protrusion and retraction behavior. Macrophages were plated at a density of 5,000 cells per dish on 35 mm glass-bottom dishes (Cellvis D35-20-1.5-N) pre-coated with 10 µg/mL fibronectin and allowed to adhere prior to imaging. Live-cell imaging was conducted using a 100x oil objective on an IX83 Olympus microscope, capturing images every 3 seconds over a 5-minute time lapse. Kymographs were generated using Olympus CellSens Count and Measure software, selecting 2–3 of the most dynamic protrusions per cell for analysis. Kymograph analysis software provided parameters such as velocity and displacement for each segment of the line profile along the cell edge. Individual segments of the line profile were then manually defined by mapping them to corresponding protruding or retracting membranes based on the movement direction of the cell edge. Additionally, the kymographs were used to manually count the number of protrusion and retraction cycles over the 5-minute imaging period.

*siRNA transfection*. Dharmacon siRNA targeting Phactr4 (Cat. L-056325) was used, along with a control non-targeting siRNA (Cat. D-001810) in a double siRNA transfection protocol. Cells were plated at a density of 150,000 cells per well in a 6-well plate and allowed to adhere overnight. For the first siRNA transfection, a final concentration of 15 nM siRNA (either Phactr4-targeting or non-targeting control) was combined with 100 µL of Opti-MEM (Thermo Fisher Cat. 31985070) and 2.5 µL of Lipofectamine™ RNAiMAX (Invitrogen Cat. 13778075) in 100 µL of Opti-MEM. The siRNA-Lipofectamine complex was incubated at room temperature for 20 minutes to allow for complex formation, and then added dropwise to the cells in complete culture media. Cells were incubated with the transfection mixture overnight, and on Day 2, the culture medium was replaced with fresh complete media. A second siRNA transfection was conducted on Day 3 using the same protocol as the first transfection. Media was again changed on Day 4. On Day 5, cells were either re-plated for downstream experiments or lysed for RNA isolation and later qRT-PCR to assess target gene knockdown.

*qRT-PCR*. Following siRNA transfection protocol, 10-15,000 cells per well were plated in 96-well plates and allowed to adhere. Cells were lysed directly in the wells to obtain RNA for analysis. cDNA synthesis and qPCR amplification were performed in a single reaction using Cells-to-CT™ 1-Step TaqMan™ Kit (Thermo Fisher, Cat. A25605) according to the manufacturer’s protocol. TaqMan™ Gene Expression Assay with FAM reporter was used to assess Phactr4 (forward: 5’-TGCCTATGTCTCAGCCTCTTC-3’, reverse: 5’-GGTCTGGGCCATAGAACTGA-3’) (Thermo Fisher, Cat. 4331182) and β-Actin (Thermo Fisher, Cat. 4386995), which served as the housekeeping control. RT-PCR reactions were conducted using QuantStudio 7 (Applied Biosystems) under the following cycling conditions: initial denaturation at 98°C for 30 seconds, followed by 40 cycles of 98°C for 10 seconds and 60°C for 30 seconds, with a final dissociation step from 65°C to 95°C in 0.5°C increments (2 seconds per step) to generate melt curves. The relative expression levels of target genes were calculated using the ΔΔCt method, normalizing to GAPDH expression.

*Co-immunoprecipitation*. Cells were resuspended in media and 4×10^5 cells were counted and pelleted at 1,000 rpm for 9 minutes. The pellet was washed once with cold 1× DPBS and resuspended in 100uL IP Lysis Buffer (50 mM Tris, pH 7.4, 1% IPEGAL, 150 mM NaCl, 1 mM EDTA, 10% glycerol, 1.5 mM MgCl₂, protease (ThermoFisher Cat. # A32953) and phosphatase (Sigma Cat # 4906845001) inhibitors. Lysates were rotated at 4°C for 15 minutes and then centrifuged at 13,000 rpm for 10 minutes. The supernatant was transferred to a new tube, and protein concentration was determined using the Precision Red protein quantification assay. If necessary, lysates were diluted to a final concentration of 1 μg/μL using IP Lysis Buffer. For input controls, 8 μg of each lysate was transferred to a separate tube and prepared for Western blot analysis. The remaining lysate was precleared using Protein G magnetic beads (DynaGreen™ ThermoFisher Cat. # 80104G). To prepare the beads, 200 μL of bead slurry was transferred to a new tube, collected using a magnetic separation rack, and washed three times with 1 mL of base lysis buffer (without inhibitors). After the final wash, beads were resuspended in 200 μL of base lysis buffer. 10 μL of the prepared Protein G magnetic bead slurry was added to each lysate sample for preclearing, followed by incubation for 30 minutes at 4°C with rotation. Beads were pelleted using a magnetic rack, and the precleared lysate was transferred to new tubes for immunoprecipitation. For immunoprecipitation, 8 μL of either Phactr4 antibody or control IgG was added to the lysates, followed by overnight incubation at 4°C with rotation. The next day, 30 μL of prepared Protein G magnetic beads was added to each immunoprecipitation sample, followed by incubation for 2 hours at 4°C with rotation. Beads were pelleted using a magnetic rack and washed three times with IP Lysis Buffer to remove non-specifically bound proteins. After the final wash, beads were resuspended in SDS sample buffer and boiled for 8 minutes at 95°C to elute immunoprecipitated proteins. Eluted samples, along with input controls, were subjected to SDS-PAGE and transferred onto PVDF membranes. Membranes were blocked in 5% milk in PBST for 1 hour at room temperature before incubation with primary antibodies overnight at 4°C. Following primary antibody incubation, membranes were washed and incubated with HRP-conjugated secondary antibodies, and protein detection was performed using chemiluminescence. Western blot images were opened in ImageJ, and the rectangular selection tool was used to define regions of interest for each band, ensuring the selection precisely encompassed the band without additional background noise. The integrated density of each protein of interest and its corresponding GAPDH band was measured, and protein expression was normalized by dividing the protein’s integrated density by that of GAPDH, with all lanes then normalized to the control lane.

*Statistical analysis*. All statistical analyses were performed using GraphPad Prism (Prism 9; GraphPad Software, San Diego, CA, USA). The statistical tests used and corresponding p-values are indicated in the figure legends. For comparisons between two groups, the Mann– Whitney test was used. For experiments involving multiple conditions and could not assume normality, ANOVAs with the Kruskal–Wallis test with Dunn’s multiple comparisons was used. Statistical significance was set at p ≤ 0.05.

## FIGURE LEGENDS

**Supplemental Figure 1.**
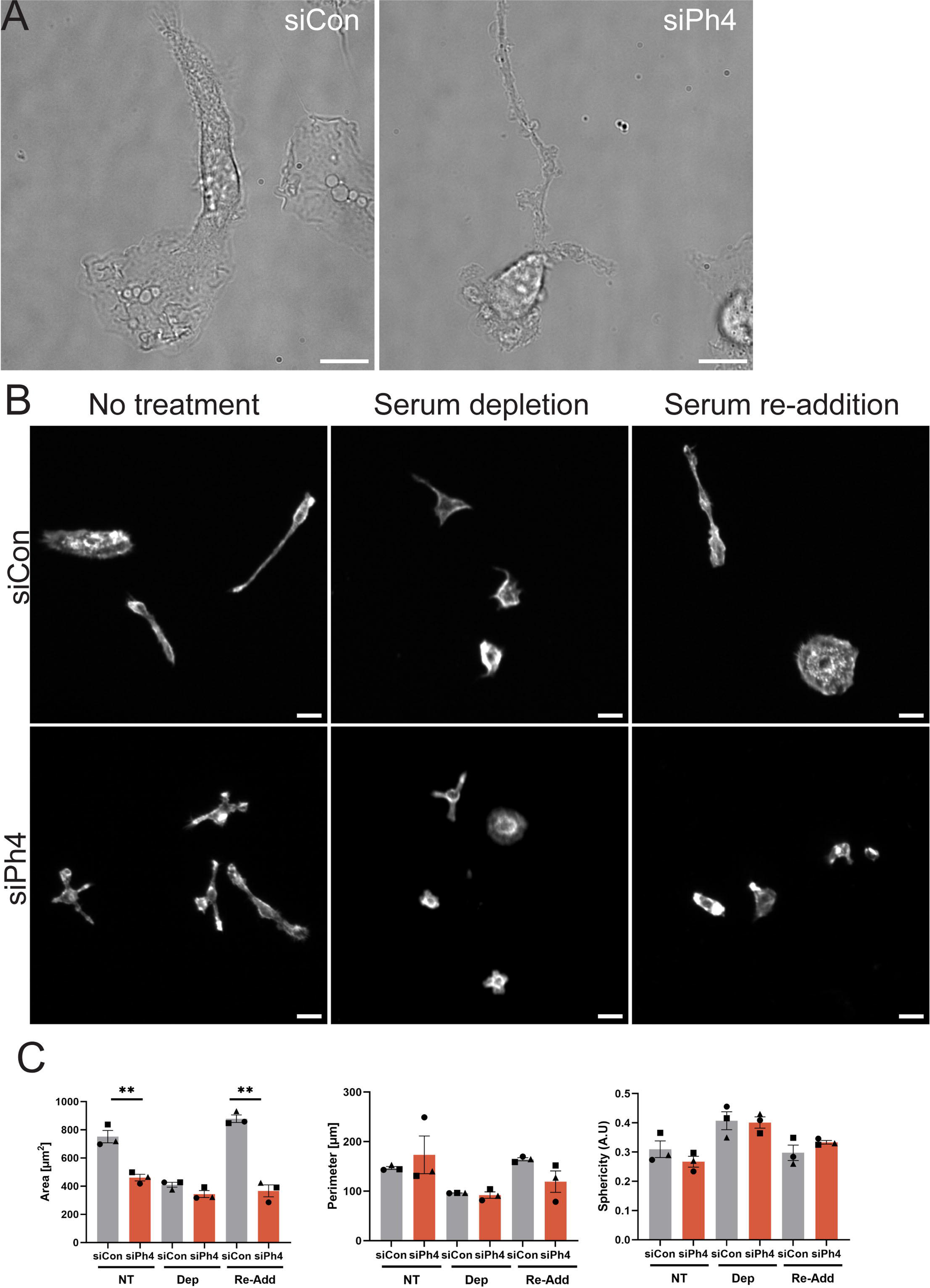
Phactr4 deletion in macrophages leads to a cell spreading defect following serum depletion and re-addition. (A) Representative differential interference contrast (DIC) images of siControl and siPhactr4 BMDMs. Cells were plated on 10 µg/mL fibronectin and allowed to spread overnight prior to imaging. Scale bar = 10 µm. (B) Representative immunofluorescence images of F-actin (gray) in siControl and siPhactr4 immortalized macrophages. Cells were either left untreated (left), subjected to serum-free media for 2 hours (middle), or serum-starved for 2 hours followed by 2 hours of serum re-addition (right) prior to fixation and staining. Scale bar = 20 µm. (C) Morphological analysis of cell area (µm²), perimeter (µm), and sphericity (arbitrary units). Automated analysis was performed using Olympus cellSens image analysis software, which masked individual cells based on F-actin fluorescence. Approximately 100 cells per condition, per experiment were analyzed, with N = 3 independent experiments. Statistical analysis was performed using Kruskal-Wallis with Dunn’s multiple comparisons test. Area (NT) siControl vs. siPhactr4 **p = 0.0091. Area (Serum Re-Addition) siControl vs. siPhactr4 **p = 0.0011.

**Supplemental Figure 2.**
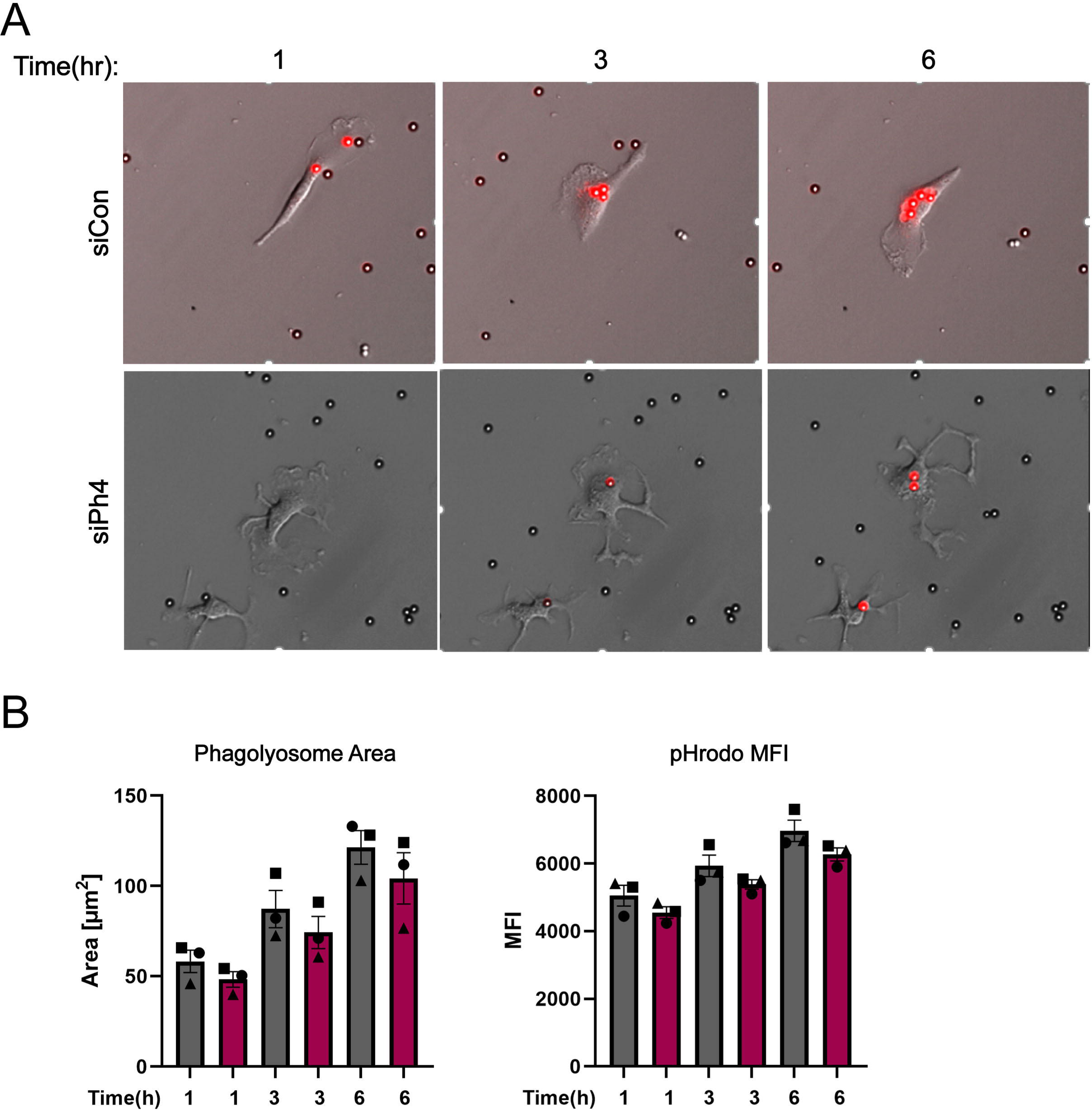
Phactr4-deficient macrophages undergo complement-mediated phagocytosis at a low rate, but demonstrate phagolysosome maturation comparable to siCon. (A) Representative differential interference contrast (DIC) images merged with the DsRed fluorescence channel of individual siControl (top) and siPhactr4 (bottom) macrophages at 1-hour, 3-hour, and 6-hour time intervals. The DsRed signal becomes fluorescent as the pHrodo-opsonized bead is internalized and enters the phagolysosome. (B) Quantification of phagolysosome area and mean fluorescence intensity (MFI) of the fluorescent signal. Phagolysosome area was quantified using Olympus CellSens image analysis software to mask the fluorescent signal at each time point of interest. The total area of each phagolysosome and the fluorescence MFI were averaged for each experiment. Data represent N = 3 independent experiments, with each experiment consisting of 10–15 fields of view and approximately 10–20 cells per frame. Statistical analysis was performed using Kruskal-Wallis with Dunn’s multiple comparisons test, with all siControl versus siPhactr4 comparisons not significant.

**Supplemental Figure 3.**
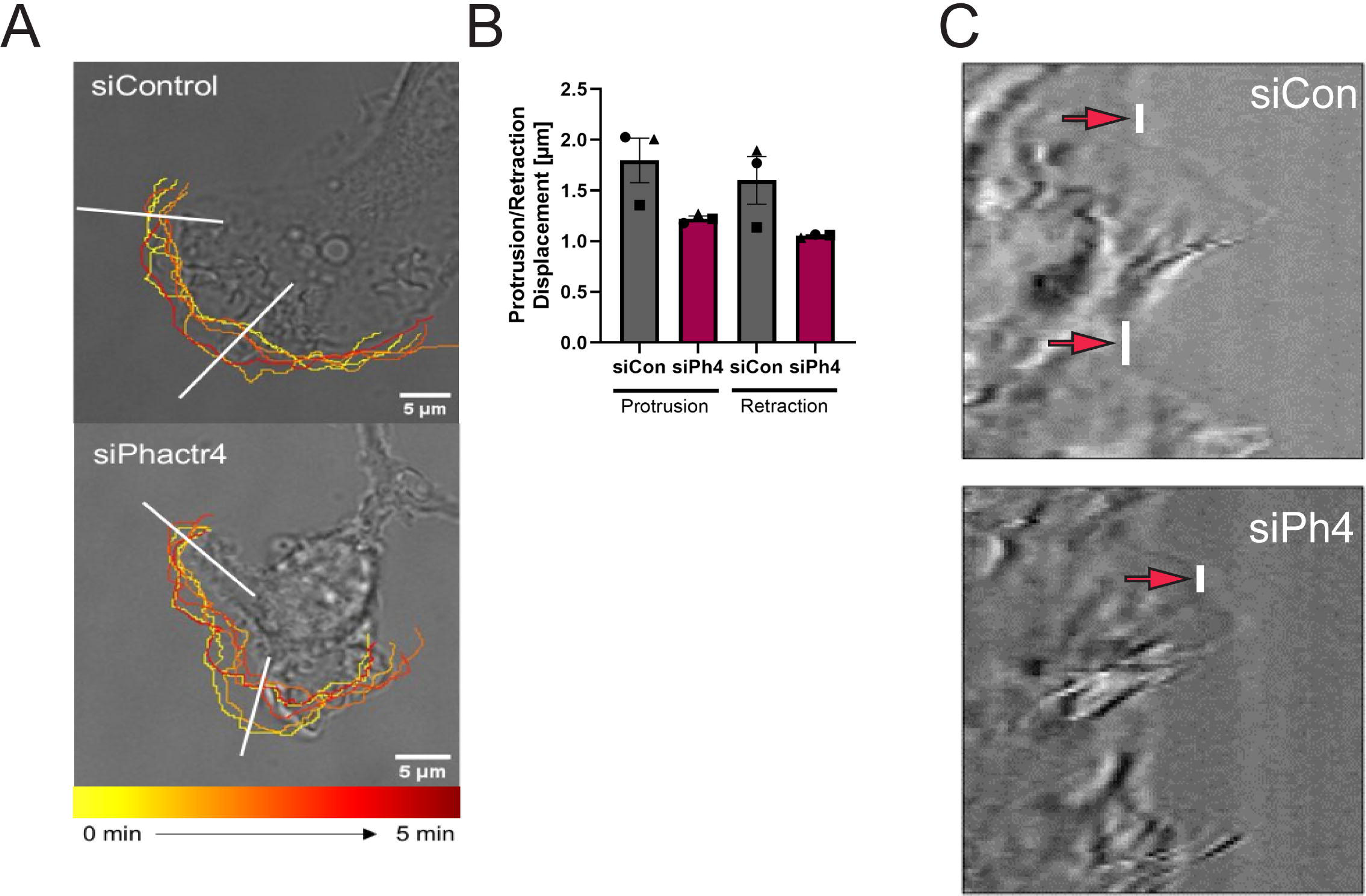
Additional kymography data for siCon and siPh4 macrophages. (A) Representative differential interference contrast (DIC) images showing a close-up of the leading edge of siControl and siPhactr4 macrophages at 100× magnification. Cells are BMDMs that underwent double siRNA transfection 7 days after isolation from bone marrow. siRNA knockdown was validated prior to imaging. Cells were plated on 10 µg/mL fibronectin and imaged 1 day after plating. Each color represents the cell edge outline at 1-minute intervals over a 5-minute period. White lines indicate 2–3 kymography lines drawn per leading edge in regions displaying visually dynamic behavior. Scale bar = 5 µm. (B) Quantification of displacement in µm of protrusion and retraction segments of siCon and siPh4 kymographs. (C) Example kymographs of siControl and siPhactr4 macrophage cell edge dynamics, highlighting stalled membrane segments. Stalled regions are indicated by white lines and red arrows.

**Supplemental Figure 4.**
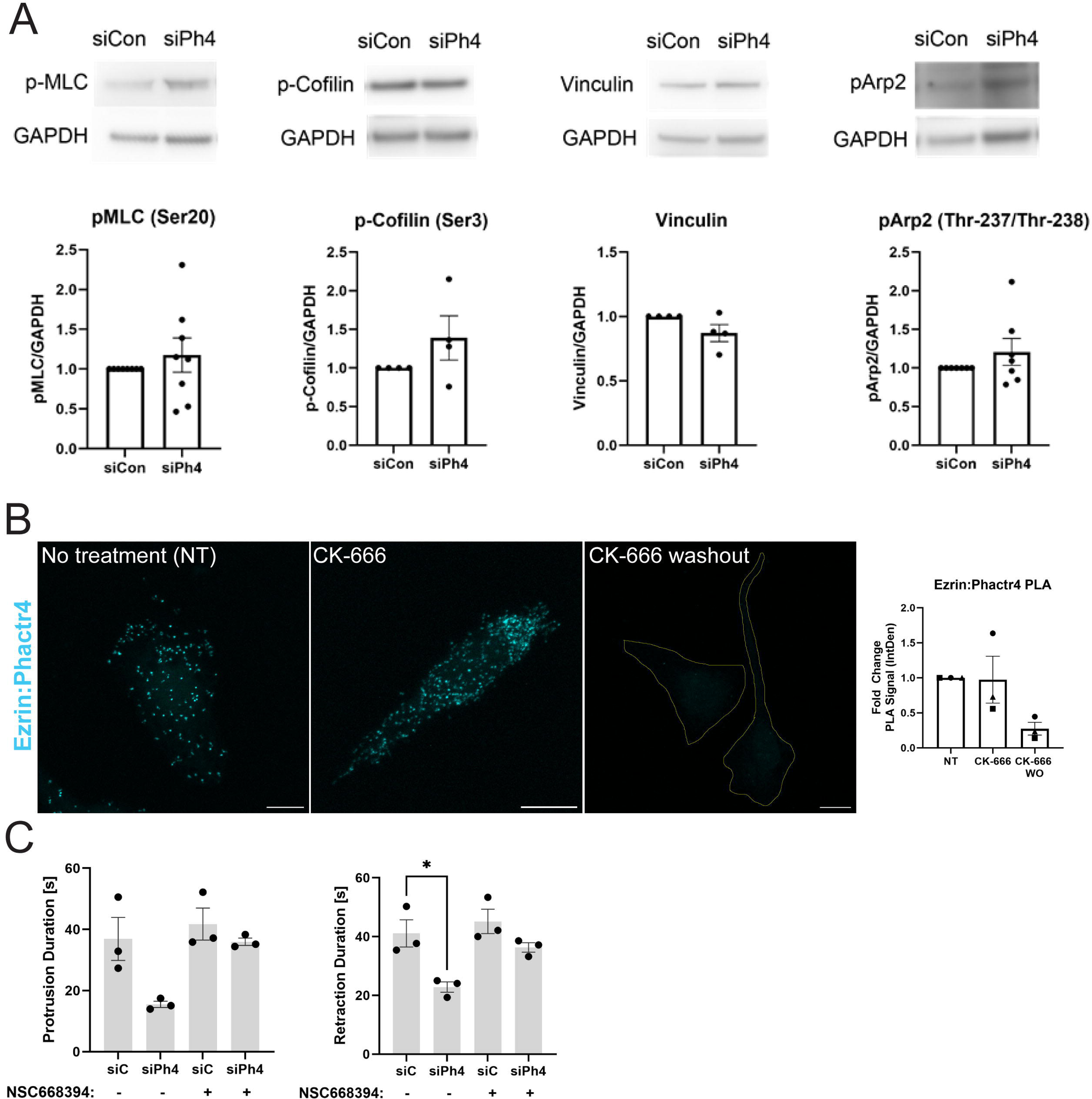
Additional phosphorylation events in siPh4 cells, rescue of siPh4 phenotype with ezrin inhibition, and supporting data on Arp2/3 complex-Phactr4 interaction. (A) Western blot analysis of (L to R) phosphorylated myosin light chain (pMLC, Ser20), phosphorylated cofilin (pCofilin, Ser3), vinculin, and phosphorylated Arp2 (pArp2, Thr-237/T-238). GAPDH was used as a loading control and for blot normalization. N = 8 for pMLC, N = 4 for pCofilin and vinculin, and N = 6 for pArp2, all independent paired siCon and siPh4 lysates. (B) Representative confocal images of proximity ligation assay (PLA) signal from the Duolink PLA assay, shown as maximum projections from a z-stack. Fluorescent puncta represent individual localizations of Ezrin-Phactr4 interactions. Cells were either untreated (left), treated with 100 µM CK-666 for 2 hours (middle), or treated with 100 µM CK-666 for 2 hours followed by a 10-minute media washout (right) before fixation and PLA assay. As signal in the washout samples was routinely low, a line was drawn around example cells in this image to denote cell volume based on companion relief contrast image. Scale bar = 10 µm for all images. Quantification of PLA fold change (right). PLA signal was normalized to the no-treatment (NT) control. Means and standard error of the mean are represented. (C) Additional kymography analysis quantification of protrusion duration and retraction duration. Statistical analysis was performed using Welch’s t-test. *\**p = 0.0444 (Retraction duration: siControl vs. siPhactr4 No Treatment).

**Supplemental Figure 5.**
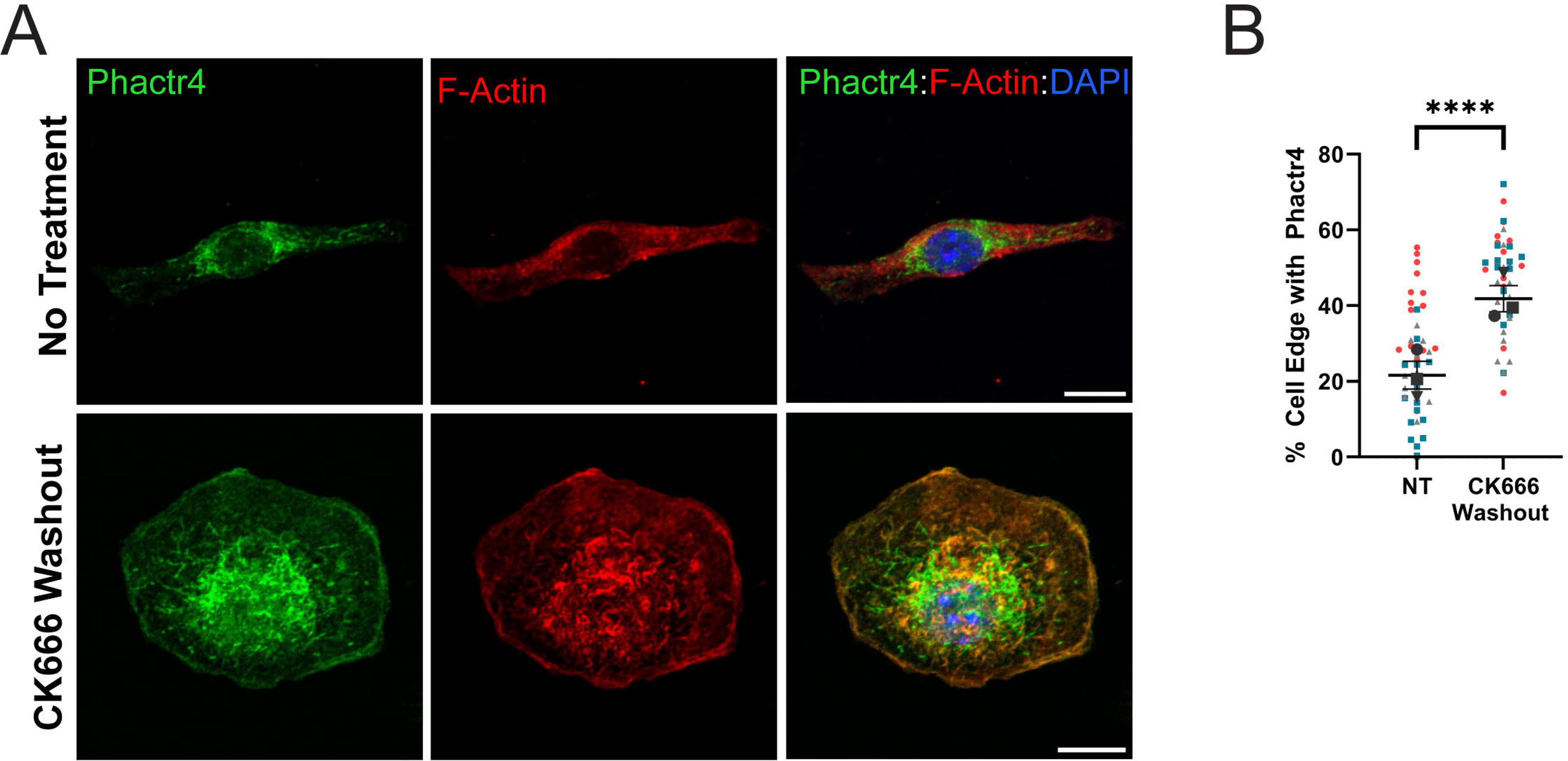
Supporting data on Arp2/3 complex-Phactr4 interaction. (A) Representative images of WT macrophages under control conditions (top) or induced spreading following CK-666 treatment and an 8-minute media washout (bottom). Scale bar = 10 µm. (B) Quantification of the percentage of the cell edge with localized Phactr4. Means and standard error of the mean are represented by black symbols, with each experiment’s mean indicated. All data points are plotted, and each experimental run is color-coded with corresponding symbols. Each experiment includes 15 cells per condition, with N = 3 independent experiments. Statistical analysis was performed using Welsh’s t-test, ****p < 0.0001.

## LIST OF ACCOMPANYING SUPPLEMENTARY MOVIES

**Supplemental Movie 1.** Time-lapse imaging of siControl (left) and siPhactr4 (right) macrophages during a random migration assay. Scale bar = 50 microns

**Supplemental Movie 2.** Time-lapse imaging of siControl (left) and siPhactr4 (right) macrophages during a phagocytosis assay. Scale bar = 50 microns

**Supplemental Movie 3.** 100x DIC imaging of siControl (left) and siPhactr4 (right) leading edges used in kymography analysis. Scale bar = 10 microns

**Supplemental Movie 4.** Time-lapse imaging of WT fibroblasts expressing GFP-Control (left) and GFP-Phactr4 (right). Scale bar = 10 microns

**Supplemental Movie 5**. 100x DIC imaging of vehicle-treated (left) and Ezrin inhibitor-treated (right) siPh4 cell leading edges used in kymography analysis. Scale bar = 10 microns

**Supplemental Movie 6.** Time-lapse imaging of WT fibroblasts expressing GFP-Phactr4 treated with DMSO (left) or CK-666 (right). Scale bar = 10 microns

## FUNDING

This work was supported by a Uniformed Services University graduate student research award (to R.M.), and by the National Institutes of Health (GM134104, to J.R.), Department of Defense (HU00012320103, to J.R.), and startup funds from the Uniformed Services University (to J.R). The Uniformed Services University of the Health Sciences (USU), 4301 Jones Bridge Rd., A1040C, Bethesda, MD 20814-4799 is the awarding and administering office.

## ACKNOWLEDGEMENTS

We thank the members of the Rotty Lab for helpful discussions during research development, and Dr. Maria Vartiainen, University of Helsinki, for her kind gift of the GFP-Phactr4 expression vector. This project is sponsored by the Uniformed Services University of the Health Sciences (USU); however, the information or content and conclusions do not necessarily represent the official position or policy of, nor should any official endorsement be inferred on the part of, USU, the Department of Defense, or the U.S. Government.

## AUTHOR CONTRIBUTIONS

R.M.: experiments and planning, data analysis, writing and editing. J.R.: Project oversight, experiments and planning, writing, editing, funding. All authors had the opportunity to review and comment on the manuscript prior to submission.

## DATA AVAILABILITY STATEMENT

All primary data will be openly available upon request.

## DECLARATION OF INTERESTS

The authors declare that they have no competing interests.

## Notes

### Competing Interest Statement

The authors have declared no competing interest.

## LITERATURE CITED

1. Mylvaganam, S., S.A. Freeman, and S. Grinstein, The cytoskeleton in phagocytosis and macropinocytosis. Current Biology, 2021. 31(10): p. R619–R632.

2. Mertz, P., V. Hentgen, G. Boursier, J. Delon, and S. Georgin-Lavialle, Current landscape of monogenic autoinflammatory actinopathies: A literature review. Autoimmunity Reviews, 2024: p. 103715.

3. Mass, E., F. Nimmerjahn, K. Kierdorf, and A. Schlitzer, Tissue-specific macrophages: how they develop and choreograph tissue biology. Nature Reviews Immunology, 2023. 23(9): p. 563–579.

4. Pollard, T.D., Regulation of actin filament assembly by Arp2/3 complex and formins. Annu. Rev. Biophys. Biomol. Struct., 2007. 36(1): p. 451–477.

5. Paul, D.S., C. Casari, C. Wu, R. Piatt, S. Pasala, R.A. Campbell, K.O. Poe, D. Ghalloussi, R.H. Lee, J.D. Rotty, et al., Deletion of the Arp2/3 complex in megakaryocytes leads to microthrombocytopenia in mice. Blood Adv, 2017. 1(18): p. 1398–1408.

6. DesMarais, V., F. Macaluso, J. Condeelis, and M. Bailly, Synergistic interaction between the Arp2/3 complex and cofilin drives stimulated lamellipod extension. Journal of cell science, 2004. 117(16): p. 3499–3510.

7. Moriyama, K., K. Iida, and I. Yahara, Phosphorylation of Ser-3 of cofilin regulates its essential function on actin. Genes to Cells, 1996. 1(1): p. 73–86.

8. Oser, M. and J. Condeelis, The cofilin activity cycle in lamellipodia and invadopodia. Journal of cellular biochemistry, 2009. 108(6): p. 1252–1262.

9. Oliver, C.J., R.T. Terry-Lorenzo, E. Elliott, W.A.C. Bloomer, S. Li, D.L. Brautigan, R.J. Colbran, and S. Shenolikar, Targeting protein phosphatase 1 (PP1) to the actin cytoskeleton: the neurabin I/PP1 complex regulates cell morphology. Molecular and cellular biology, 2002. 22(13): p. 4690–4701.

10. Ambach, A., J. Saunus, M. Konstandin, S. Wesselborg, S.C. Meuer, and Y. Samstag, The serine phosphatases PP1 and PP2A associate with and activate the actin-binding protein cofilin in human T lymphocytes. European journal of immunology, 2000. 30(12): p. 3422–3431.

11. Martin-Granados, C., A.R. Prescott, N. Van Dessel, A. Van Eynde, M. Arocena, I.P. Klaska, J. Görnemann, M. Beullens, M. Bollen, and J.V. Forrester, A role for PP1/NIPP1 in steering migration of human cancer cells. PloS one, 2012. 7(7): p. e40769.

12. Heroes, E., B. Lesage, J. Görnemann, M. Beullens, L. Van Meervelt, and M. Bollen, The PP1 binding code: a molecular-lego strategy that governs specificity. The FEBS journal, 2013. 280(2): p. 584–595.

13. Zhang, Y., T.-H. Kim, and L. Niswander, Phactr4 regulates directional migration of enteric neural crest through PP1, integrin signaling, and cofilin activity. Genes & development, 2012. 26(1): p. 69–81.

14. Bach, I., The LIM domain: regulation by association. Mech Dev, 2000. 91(1-2): p. 5–17.

15. Solimini, N.L., A.C. Liang, C. Xu, N.N. Pavlova, Q. Xu, T. Davoli, M.Z. Li, K.-K. Wong, and S.J. Elledge, STOP gene Phactr4 is a tumor suppressor. Proceedings of the National Academy of Sciences, 2013. 110(5): p. E407–E414.

16. Cao, F., M. Liu, Q. Zhang, and R. Hao, PHACTR4 regulates proliferation, migration and invasion of human hepatocellular carcinoma by inhibiting IL-6/Stat3 pathway. Eur Rev Med Pharmacol Sci, 2016. 20(16): p. 3392–3399.

17. Pandharipande, P.P., R.D. Sanders, T.D. Girard, S. McGrane, J.L. Thompson, A.K. Shintani, D.L. Herr, M. Maze, E.W. Ely, and M. investigators, Effect of dexmedetomidine versus lorazepam on outcome in patients with sepsis: an a priori-designed analysis of the MENDS randomized controlled trial. Crit Care, 2010. 14(2): p. R38.

18. Lamb, R.F., B.W. Ozanne, C. Roy, L. McGarry, C. Stipp, P. Mangeat, and D.G. Jay, Essential functions of ezrin in maintenance of cell shape and lamellipodial extension in normal and transformed fibroblasts. Current Biology, 1997. 7(9): p. 682–688.

19. Rouven Brückner, B., A. Pietuch, S. Nehls, J. Rother, and A. Janshoff, Ezrin is a major regulator of membrane tension in epithelial cells. Scientific reports, 2015. 5(1): p. 14700.

20. Fievet, B.T., A. Gautreau, C. Roy, L. Del Maestro, P. Mangeat, D. Louvard, and M. Arpin, Phosphoinositide binding and phosphorylation act sequentially in the activation mechanism of ezrin. The Journal of cell biology, 2004. 164(5): p. 653–659.

21. Matsui, T., M. Maeda, Y. Doi, S. Yonemura, M. Amano, K. Kaibuchi, S. Tsukita, and S. Tsukita, Rho-kinase phosphorylates COOH-terminal threonines of ezrin/radixin/moesin (ERM) proteins and regulates their head-to-tail association. J Cell Biol, 1998. 140(3): p. 647–57.

22. Proudfit, A., N. Bhunia, D. Pore, Y. Parker, D. Lindner, and N. Gupta, Pharmacologic Inhibition of Ezrin-Radixin-Moesin Phosphorylation is a Novel Therapeutic Strategy in Rhabdomyosarcoma. Sarcoma, 2020. 2020(1): p. 9010496.

23. Zhang, X., L.R. Flores, M.C. Keeling, K. Sliogeryte, and N. Gavara, Ezrin phosphorylation at T567 modulates cell migration, mechanical properties, and cytoskeletal organization. International journal of molecular sciences, 2020. 21(2): p. 435.

24. Welf, E.S., C.E. Miles, J. Huh, E. Sapoznik, J. Chi, M.K. Driscoll, T. Isogai, J. Noh, A.D. Weems, and T. Pohlkamp, Actin-membrane release initiates cell protrusions. Developmental cell, 2020. 55(6): p. 723–736. e8.

25. Lawson, C.D. and K. Burridge, The on-off relationship of Rho and Rac during integrin-mediated adhesion and cell migration. Small GTPases, 2014. 5: p. e27958.

26. Dobson, L., W.B. Barrell, Z. Seraj, S. Lynham, S.Y. Wu, M. Krause, and K.J. Liu, GSK3 and lamellipodin balance lamellipodial protrusions and focal adhesion maturation in mouse neural crest migration. Cell Rep, 2023. 42(9): p. 113030.

27. Michael, M., A. Vehlow, C. Navarro, and M. Krause, c-Abl, Lamellipodin, and Ena/VASP proteins cooperate in dorsal ruffling of fibroblasts and axonal morphogenesis. Curr Biol, 2010. 20(9): p. 783–91.

28. Benz, P.M., C. Blume, S. Seifert, S. Wilhelm, J. Waschke, K. Schuh, F. Gertler, T. Munzel, and T. Renne, Differential VASP phosphorylation controls remodeling of the actin cytoskeleton. J Cell Sci, 2009. 122(Pt 21): p. 3954–65.

29. Lebrand, C., E.W. Dent, G.A. Strasser, L.M. Lanier, M. Krause, T.M. Svitkina, G.G. Borisy, and F.B. Gertler, Critical role of Ena/VASP proteins for filopodia formation in neurons and in function downstream of netrin-1. Neuron, 2004. 42(1): p. 37–49.

30. Singh, S.P., P.A. Thomason, and R.H. Insall, Extracellular Signalling Modulates Scar/WAVE Complex Activity through Abi Phosphorylation. Cells, 2021. 10(12).

31. Singh, S.P., P.A. Thomason, S. Lilla, M. Schaks, Q. Tang, B.L. Goode, L.M. Machesky, K. Rottner, and R.H. Insall, Cell-substrate adhesion drives Scar/WAVE activation and phosphorylation by a Ste20-family kinase, which controls pseudopod lifetime. PLoS Biol, 2020. 18(8): p. e3000774.

32. Arber, S., F.A. Barbayannis, H. Hanser, C. Schneider, C.A. Stanyon, O. Bernard, and P. Caroni, Regulation of actin dynamics through phosphorylation of cofilin by LIM-kinase. Nature, 1998. 393(6687): p. 805–9.

33. Yang, N., O. Higuchi, K. Ohashi, K. Nagata, A. Wada, K. Kangawa, E. Nishida, and K. Mizuno, Cofilin phosphorylation by LIM-kinase 1 and its role in Rac-mediated actin reorganization. Nature, 1998. 393(6687): p. 809–12.

34. LeClaire III, L.L., M. Baumgartner, J.H. Iwasa, R.D. Mullins, and D.L. Barber, Phosphorylation of the Arp2/3 complex is necessary to nucleate actin filaments. The Journal of cell biology, 2008. 182(4): p. 647–654.

35. LeClaire, L.L., M. Rana, M. Baumgartner, and D.L. Barber, The Nck-interacting kinase NIK increases Arp2/3 complex activity by phosphorylating the Arp2 subunit. Journal of Cell Biology, 2015. 208(2): p. 161–170.

36. Cai, L., A.M. Makhov, D.A. Schafer, and J.E. Bear, Coronin 1B antagonizes cortactin and remodels Arp2/3-containing actin branches in lamellipodia. Cell, 2008. 134(5): p. 828–42.

37. Cai, L., T.W. Marshall, A.C. Uetrecht, D.A. Schafer, and J.E. Bear, Coronin 1B coordinates Arp2/3 complex and cofilin activities at the leading edge. Cell, 2007. 128(5): p. 915–29.

38. Gandhi, M., B.A. Smith, M. Bovellan, V. Paavilainen, K. Daugherty-Clarke, J. Gelles, P. Lappalainen, and B.L. Goode, GMF is a cofilin homolog that binds Arp2/3 complex to stimulate filament debranching and inhibit actin nucleation. Curr Biol, 2010. 20(9): p. 861–7.

39. Dang, I., R. Gorelik, C. Sousa-Blin, E. Derivery, C. Guerin, J. Linkner, M. Nemethova, J.G. Dumortier, F.A. Giger, T.A. Chipysheva, et al., Inhibitory signalling to the Arp2/3 complex steers cell migration. Nature, 2013. 503(7475): p. 281–4.

40. Cai, L., N. Holoweckyj, M.D. Schaller, and J.E. Bear, Phosphorylation of coronin 1B by protein kinase C regulates interaction with Arp2/3 and cell motility. J Biol Chem, 2005. 280(36): p. 31913–23.

41. Zhang, Y. and L. Niswander, Phactr4: a new integrin modulator required for directional migration of enteric neural crest cells. Cell adhesion & migration, 2012. 6(5): p. 419–423.

42. Nishimura, T., T. Yamaguchi, K. Kato, M. Yoshizawa, Y.-i. Nabeshima, S. Ohno, M. Hoshino, and K. Kaibuchi, PAR-6–PAR-3 mediates Cdc42-induced Rac activation through the Rac GEFs STEF/Tiam1. Nature cell biology, 2005. 7(3): p. 270–277.

43. Nishimura, T. and K. Kaibuchi, Numb controls integrin endocytosis for directional cell migration with aPKC and PAR-3. Developmental cell, 2007. 13(1): p. 15–28.

44. Iden, S. and J.G. Collard, Crosstalk between small GTPases and polarity proteins in cell polarization. Nature reviews Molecular cell biology, 2008. 9(11): p. 846–859.

45. Traweger, A., G. Wiggin, L. Taylor, S.A. Tate, P. Metalnikov, and T. Pawson, Protein phosphatase 1 regulates the phosphorylation state of the polarity scaffold Par-3. Proceedings of the National Academy of Sciences, 2008. 105(30): p. 10402–10407.

46. Martin, T.A., G. Harrison, R.E. Mansel, and W.G. Jiang, The role of the CD44/ezrin complex in cancer metastasis. Critical reviews in oncology/hematology, 2003. 46(2): p. 165–186.

47. Meng, Y., Z. Lu, S. Yu, Q. Zhang, Y. Ma, and J. Chen, Ezrin promotes invasion and metastasis of pancreatic cancer cells. Journal of translational medicine, 2010. 8: p. 1–14.

48. Cao, F., M. Liu, Q.Z. Zhang, and R. Hao, PHACTR4 regulates proliferation, migration and invasion of human hepatocellular carcinoma by inhibiting IL-6/Stat3 pathway. Eur Rev Med Pharmacol Sci, 2016. 20(16): p. 3392–9.

49. Barik, G.K., O. Sahay, D. Paul, and M.K. Santra, Ezrin gone rogue in cancer progression and metastasis: An enticing therapeutic target. Biochimica et Biophysica Acta (BBA)-Reviews on Cancer, 2022. 1877(4): p. 188753.

50. Shao, S., H. Miao, and W. Ma, Unraveling the enigma of tumor-associated macrophages: challenges, innovations, and the path to therapeutic breakthroughs. Front Immunol, 2023. 14: p. 1295684.

51. Zha, H., X. Wang, Y. Zhu, D. Chen, X. Han, F. Yang, J. Gao, C. Hu, C. Shu, and Y. Feng, Intracellular activation of complement C3 leads to PD-L1 antibody treatment resistance by modulating tumor-associated macrophages. Cancer immunology research, 2019. 7(2): p. 193–207.

52. Li, Y., X. Chen, T. Lan, W. Wang, C. Wang, M. Chang, Z. Yu, and S. Yu, Targeting Phactr4 to rescue chronic stress-induced depression-like behavior in rats via regulating neuroinflammation and neuroplasticity. International Journal of Biological Macromolecules, 2024. 273: p. 132854.

53. Rotty, J.D., H.E. Brighton, S.L. Craig, S.B. Asokan, N. Cheng, J.P. Ting, and J.E. Bear, Arp2/3 Complex Is Required for Macrophage Integrin Functions but Is Dispensable for FcR Phagocytosis and In Vivo Motility. Dev Cell, 2017. 42(5): p. 498–513 e6.

54. Huet, G., E.K. Rajakyla, T. Viita, K.P. Skarp, M. Crivaro, J. Dopie, and M.K. Vartiainen, Actin-regulated feedback loop based on Phactr4, PP1 and cofilin maintains the actin monomer pool. J Cell Sci, 2013. 126(Pt 2): p. 497–507.

55. Stinson, M.W., S. Liu, A.J. Laurenson, and J.D. Rotty, Macrophage migration is differentially regulated by fibronectin and laminin through altered adhesion and myosin II localization. Mol Biol Cell, 2024. 35(2): p. ar22.

